# Foundation model reveals the shared organization of transcription and topologically associating domains

**DOI:** 10.1101/2025.03.31.646349

**Authors:** Huan Liang, Bonnie Berger, Rohit Singh

## Abstract

The three-dimensional organization of chromatin into topologically associating domains (TADs) may impact gene regulation by bringing distant genes into contact. However, studies of TADs’ function and their influence on transcription have been constrained by ambiguities in TAD boundary definitions and challenges in directly measuring their regulatory effects. We overcome these limitations by developing species-level consensus TAD maps for human and mouse using a bag-of-genes approach that exposes emergent regulatory structure. To quantify TAD-mediated gene relationships, we leverage a generative AI foundation model computed from 33 million transcriptomes to define a contextual similarity metric, revealing higher-order gene relationships elusive to co-expression analysis. We find that TADs are regions of elevated gene co-regulation, and our analytical framework directly leads to testable hypotheses about chromatin organization across cellular states. We discover that the TAD-linked enhancement of transcriptional context is strongest in early developmental stages and systematically declines with aging. Investigation of cancer cells show distinct patterns of TAD usage that shift with chemotherapy treatment. Enhancing our understanding of cellular plasticity in differentiation and disease, these findings suggest that chromatin organization may act through probabilistic mechanisms rather than deterministic rules.

**Software availability:** https://singhlab.net/tadmap

## Introduction

Cells must precisely orchestrate the activity of hundreds of genes to perform even basic functions. This is fundamentally a physical problem, and evolution has found distinct solutions. Bacteria use operons to align co-regulated genes along the genome^1^; mammals fold DNA within the crowded nucleus, employing dynamic three-dimensional chromatin architecture that brings distant genes into contact^2, 3^. Topologically associating domains (TADs), unveiled by diverse chromatin conformation capture technologies^4–8^, are key components of this hierarchical architecture^2, 9–11^—regions where DNA frequently contacts within the domain but rarely crosses it. These domains create distinct neighborhoods within the genome^12–14^ in which the transcriptional machinery operates.

TADs present a paradox. Although their importance is clear, their function remains disputed. Do they coordinate or insulate gene activity? How do they relate to the genome’s linear organization? Even these basic questions remain unresolved^15^. This uncertainty stems from technical challenges: TAD boundaries vary between cell types and individual cells; studies of CTCF, a protein central to TAD formation, have yielded inconsistent results^16–21^. The complexity of quantifying gene co-transcription exacerbates the challenge. Bulk RNA-seq datasets have limited sample sizes, while single-cell RNA-seq introduces technical noise and data sparsity. Some gene-dense TADs, such as those containing olfactory receptor genes, show minimal co-expression^22, 23^. This raises a central question: are there general principles governing how TAD organization relates to transcription, or do these relationships vary too unpredictably with context to generalize? To answer this question, we zoom out from individual cell-type analyses to the species level.

At the species level, a striking regularity emerges. Genes sharing a TAD show 20.5% higher transcriptional similarity than genes at equivalent distances outside TADs, an enhancement that persists multiplicatively across all distance scales and holds across diverse cell types. This pattern had previously been difficult to detect. Precise TAD boundaries vary across cell types and whether TADs represent conserved regulatory units or are artifacts of chromatin conformation remains debated^24^. For measuring co-transcription, the standard metric, pairwise Pearson expression correlation, struggles with sparse single-cell data and misses genes that serve equivalent roles without being expressed together. Our species-level analysis, integrating Hi-C data over dozens of cell types with tens of millions of single-cell transcriptomes, exposes a consistent quantitative relationship: transcriptional coordination within TADs is systematically elevated in ways genomic proximity alone cannot explain.

We develop two conceptual advances that reveal this pattern. The *TAD Map* abstracts chromatin structure to the species level, representing each TAD simply as its contained genes, a “bag of genes” model (Fig. 1A) inspired by set-based representations in language and protein modeling^25^. This abstraction tolerates boundary variation^26–30^ and enables genome-wide analysis without requiring cell type–specific Hi-C data. *Contextual transcriptional similarity* (CTS) applies single-cell foundation models to a new purpose: detecting gene relationships that co-expression misses. Derived from scGPT, a foundation model trained on 33 million cells, CTS captures whether genes occupy similar roles within cellular programs. Consider genes as Lego blocks, each fitting into specific transcriptional roles defined by other genes. Many genes have unique shapes, but certain families, such as olfactory receptors and protocadherins, act like blocks of identical shape but different colors. Naive co-expression misses relationships among such genes; CTS exposes their shared context.

**Figure 1:**
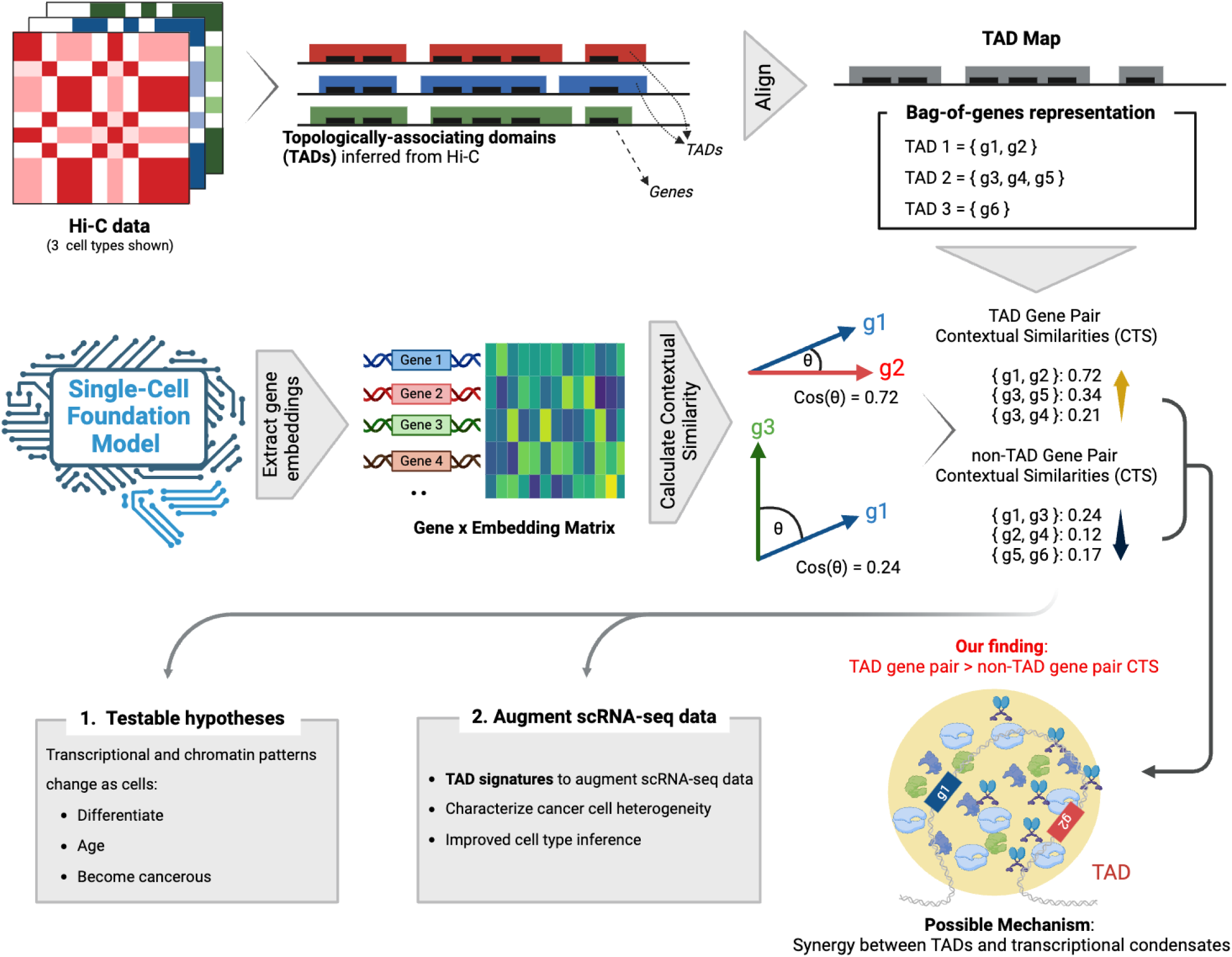
Overview. Topologically associating domains (TADs) are a key folding unit of the chromatin. While TADs are currently inferred separately for each cell type, TAD architectures have been observed to have good agreement across cell types. We find this agreement to be even stronger when TADs are represented simply as sets of genes, suggesting a bag-of-genes representation would have species-wide applicability. Applying maximum likelihood estimation, we compute a consensus TAD scaffold from Hi-C data and the corresponding bag-of-genes representation (the *TAD Map*, Fig. 2). The TAD Map enables us to “impute” the high-level chromatin structure in each cell/tissue. We demonstrate that the genome partitioning implied by the TAD scaffold agrees with functional genomic data across a variety of cell/tissue types. (ii) We leverage gene representations from single-cell foundation models, pretrained with more than 33 million cells, to derive contextual transcriptional similarities (CTS) between gene pairs. These measures capture co-regulation beyond simple expression-correlation patterns between two genes (Fig. 3). (iii) We find CTS in TAD gene pairs significantly exceeds that in non-TAD gene pairs and hypothesize that a synergy between TADs and transcriptional condensates may explain it (Fig. 4). The TAD Map framework enables novel, testable hypothesis: We examine how TAD usage and CTS change as cells develop and age, or become cancerous (Fig. 4). (iv) We introduce *TAD signatures*, a probabilistic model of TAD activation inferred from single-cell RNA-seq (scRNA-seq) readouts, showing how they facilitate greater accuracy and robustness in downstream scRNA-seq analyses (Fig. 5).

Beyond this core finding, our framework generates testable predictions about chromatin organization during development, aging, and cancer, and we evaluate these predictions using diverse single-cell datasets. We also introduce TAD signatures, bringing species-level chromatin structure to single-cell RNA-seq analysis and improving interpretation of individual transcriptomes.

## Results

### Consensus TAD Maps for human and mouse

The genome’s three-dimensional organization presents a substantial consistency across cell types^26–32^. To quantify this consistency, we analyzed TAD definitions from seven human and four mouse cell types using the TADKB database^33^ and the Directionality Index technique^26^. The TADKB database provided harmonized TAD calls across diverse data sources (**Methods**). This analysis revealed high conservation (**Fig. 2A**). In humans, 92.6% of TAD boundaries matched those in at least one other cell type (within 50 kb), though only 31.7% matched across *all* cell types. Mouse showed lower conservation, with corresponding scores of 69.9% and 13.4%. We extended this analysis using data from the 4D Nucleome project^34^, incorporating ten human and six mouse additional cell types. Analyzing with the Directionality Index as well as the Insulation Score methods for TAD-calling^35^, we found broadly consistent results (**Fig. S1**).

**Figure 2:**
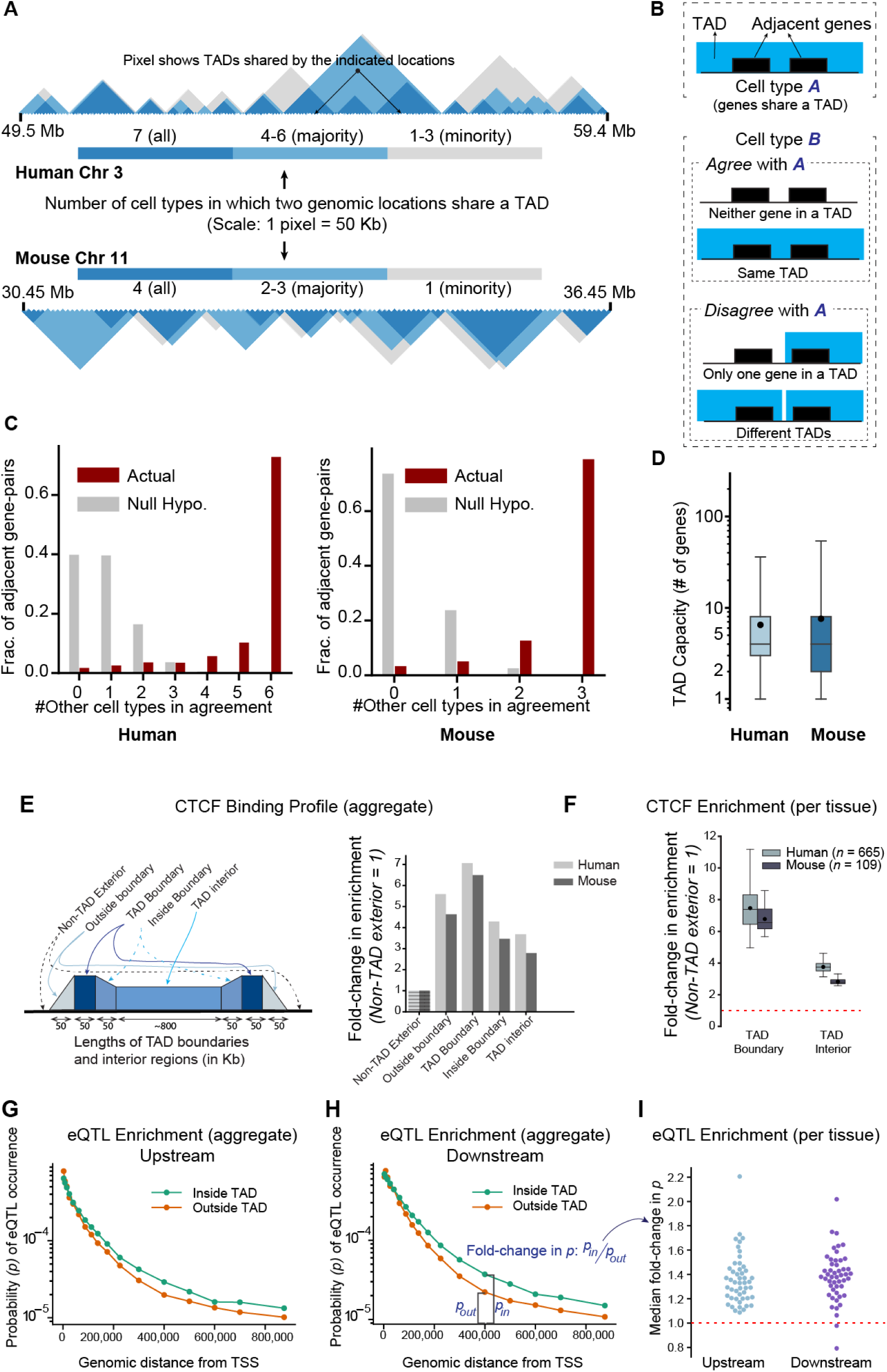
Agreement between cell-type specific TAD architectures and the constructed consensus TAD scaffold. **A**) On representative segments of the human and mouse genomes, overlap of cell-type specific TAD architectures (7 cell types for human, 4 four mouse) are shown. Genomic positional range (in Mb) is shown on the horizontal axis and the number of cell types in which a pair of genomic loci co-occupy a TAD is indicated by the shaded triangles. In each species, the majority of the cell-type specific TAD architectures are in agreement on most genomic loci pairs; in many cases, *all* TAD architectures are in agreement. **B**) Schematic for the statistical test to evaluate if two bag-of-genes TAD representations are identical: given an adjacently-located gene pair that shares a TAD in at least one cell type’s TAD architecture, we mark other cell type(s) to be in agreement if the gene pair is not separated by a TAD boundary. **C**) For each adjacent gene pair that shares a TAD in at least one cell type, the number of other cell types where the genes share a TAD. Under this test, 72.7% of adjacent gene pairs in human (78.9% in mouse) show complete agreement across all cell-type specific TAD architectures. The expected count under a simple null hypothesis (Methods) is below 1 in both species. **D**) Our consensus TAD scaffolds for each of human and mouse, computed by maximum likelihood estimation, yield 3036 (3181) TADs with an average length of 879 kb (845 kb) in the human (mouse) genome, with the median TAD in both species containing 4 genes. In this figure, the box represents the 25-75^*th*^ percentile range, and the whiskers represent 1-99^*th*^ percentile range. **E, F**) Analysis of 665 human and 109 mouse CTCF ChIP-seq assays (ENCODE) revealed significant enrichment of CTCF binding at TAD boundaries and within TADs. Binding strength declined on either side of the predicted boundaries, supporting the accuracy of the scaffold. F) In individual tissues, CTCF enrichment at TAD boundaries was highly significant (Bonferroni-corrected *p* < 10^−50^, one-sided binomial test). The box in the figure represents the 25-75^*th*^ percentile, while the whiskers show the 1-99^*th*^ percentile range. **G, H, I**) To assess the functional relevance of the TAD scaffold, we compared eQTL prevalence inside and outside TADs using single-tissue data from GTEx v8. Loci within the same TAD as a gene’s transcription start site (TSS) were associated with significantly higher eQTL frequencies (*p* = 0.0001 for both upstream and downstream loci, one-sided Wilcoxon rank sum test). This trend held true for most tissues, demonstrating the scaffold’s informativeness across diverse tissue types.

While the boundaries of TADs vary, we reason that their gene memberships may be more stable. We therefore represent each TAD simply as its set of protein-coding genes, a “bag-of-genes” abstraction we call the TAD Map.

This representation tolerates minor boundary shifts that don’t alter gene membership while retaining the features needed for statistical analysis of gene regulation. The bag-of-genes approach reveals strong consistency across cell types. We assessed this by examining adjacent gene pairs that share a TAD in at least one cell type (**Fig. 2B**). In humans, 72.8% of these pairs remain grouped together across *all* cell types, rising to 92.3% when considering majority agreement. Mouse shows similar patterns (78.9% and 91.6% respectively). This consistency significantly exceeds expectations under a null model of independent boundary dispersion (one-sided binomial test, *p* = 7.3 × 10^−75^ for human, 1.7 × 10^−4^ for mouse; **Fig. 2C, Methods**). We extended the analysis to gene triplets, which yielded similar findings (**Fig. S2**).

Consensus TAD “scaffolds” organize the genome into gene groupings preserved across cell types. We identified optimal TAD boundaries by partitioning chromosomes into 50 kb segments and applying dynamic programming to find non-overlapping intervals best supported by TADKB data (**Methods**). The human scaffold comprises 3,036 TADs (mean length 879 kb, median 4 protein-coding genes per TAD), while the mouse scaffold contains 3,181 TADs (mean length 845 kb, median 4 protein-coding genes per TAD) (**Fig. 2D**). Together, human and mouse TADs cover approximately 86% and 92% of their respective genomes. From these scaffolds, we derive the complete TAD Map by associating each TAD with all its genes, including those spanning multiple TADs (**Discussion**).

#### TAD Map agrees with CTCF ChIP-seq data

CTCF binding validates the TAD Map across hundreds of cell types lacking Hi-C data. CTCF enrichment at TAD boundaries^36, 37^ (and sometimes inside them^3, 38^) is well-established, but prior studies were limited to cell types with available Hi-C. We sourced CTCF ChIP-seq data from the ENCODE database^39^, comprising 281 human and 28 mouse studies (665 and 109 replicates, respectively). These studies include many cell types where TAD–CTCF concordance could not be previously studied due to the lack of Hi-C data. We find that in both human and mouse, CTCF binding occurs most frequently at TAD boundaries and remains significantly higher inside TADs than in the general genomic background (**Methods**, **Fig. 2E**). This pattern persists across diverse cell types and disease states (**Fig. 2F**), supporting the broad applicability of the TAD Map.

#### Genetic variants within TADs show elevated regulatory effects on nearby genes

We analyzed expression quantitative trait loci (eQTLs), spanning all 49 tissues covered in GTEx v8^40^. For each gene, we estimated the probability *p*(*d*) that a variant at distance *d* from its transcription start site (TSS) functions as an eQTL (**Methods**). As reported previously^41, 42^, this probability declines with distance. However, we find this decline is significantly slower inside TADs (**Fig. 2G**). Beyond the immediate vicinity of genes (*d* > 30 kb), variants within TADs show higher probability of being eQTLs (**Fig. 2H**), with median fold changes of 1.27 and 1.37 for upstream and downstream regions, respectively (*p*-values < 5 × 10^−4^, Wilcoxon rank sum test). These results, consistent across nearly all tissues, further validate our TAD Map, indicating that TAD membership influences the regulatory reach of genetic variants.

### Single-cell foundation models characterize the transcriptional context of genes

Co-expression analysis misses a fundamental aspect of gene relationships: functional substitutability. In olfactory neurons, only one of several hundred receptors is typically expressed, yet all serve equivalent roles in specifying neuronal identity^22^. Protocadherin genes show similar patterns^43^. These genes often reside in the same TADs, suggesting TAD-based gene groupings imply functional relationships that traditional correlation-based measures are unable to fully capture.

Single-cell foundation models (scFMs) offer an alternative. These models integrate tens of millions of transcriptomes to learn gene representations that reflect broader functional context^44–46^. We leverage scGPT, trained on 33 million human cells, to define “Contextual Transcriptional Similarity” (CTS): the Pearson correlation between two genes’ embedding vectors. Unlike co-expression, which requires observing genes active together in the same cells, CTS characterizes a gene’s role through its relationships with all other genes across diverse cellular states. Two genes can thus show high CTS despite rarely co-expressing—capturing substitutability that correlation-based metrics miss.

CTS captures greater variation in gene relationships than co-expression. Analyzing 418,425 protein-coding gene pairs (less than 2 megabases apart on the same chromosome) across four atlas-scale scRNA-seq datasets from CELLxGENE^47–51^, we found that CTS shows significantly higher variation than co-expression (variances of 0.010 vs. 0.002 after Fisher’s transformation, *p* < 10^−15^, one-sided F test; **Fig. 3A-B**). The compressed range of co-expression values likely reflects single-cell data sparsity, where many transcripts per cell are reported as zero.

**Figure 3:**
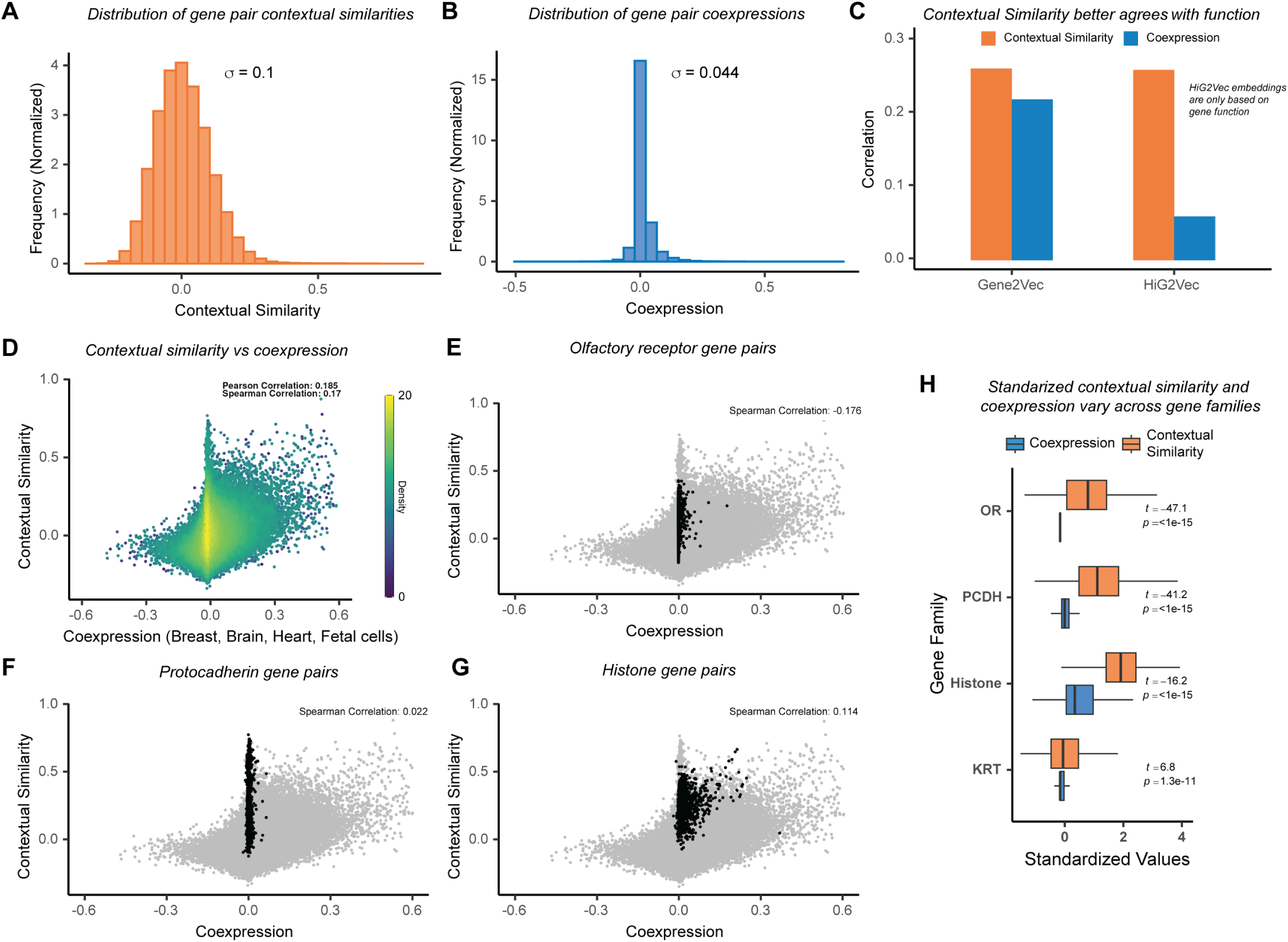
Contextual similarity offers a more holistic perspective in evaluating gene pair transcriptional patterns. A, **B**) Using Pearson correlations of pre-trained scGPT gene embeddings, we calculated contextual transcriptional similarities (CTS) for all gene pairs on the same chromosome within 2 megabases. Gene pair co-expression was similarly computed using tissue-specific single-cell data. CTS proved to be a more diverse and statistically powerful metric for evaluating gene interactions than co-expression. **C**) We compared CTS and co-expression against similarities derived from two popular gene embedding methods: Gene2Vec and HiG2Vec. Gene2Vec incorporates both gene expression and functional annotations, while HiG2Vec is based solely on functional annotations. CTS correlates more strongly than co-expression with both measures, particularly with the gene ontology-based HiG2Vec. This suggests CTS can effectively distill raw expression data to capture gene function in its embeddings. **D**) CTS and co-expression show positive correlation across all gene pairs, confirming that both metrics capture shared aspects of gene expression patterns. **E, F, G**) Analysis of three gene families—olfactory receptors (ORs), protocadherins (PCDHs), and histones—reveals CTS’s ability to capture functional relationships beyond co-expression. ORs and PCDHs show minimal co-expression by design (ORs are expressed one gene at a time, PCDHs stochastically), yet CTS identifies their shared functional roles. In contrast, histones, which are coordinately expressed, show high values in both metrics. **H**) Quantification across gene families shows CTS captures a broader range of relationships than co-expression, with the strongest signal in histones, followed by PCDHs and ORs, matching their known biological relationships. Additional analysis of keratin genes further supports this pattern (Fig. S2).

CTS correlates more strongly with functional annotations than co-expression does. While CTS and co-expression show broad agreement (**Fig. 3D**; Pearson correlation 0.185, *p* < 10^−15^), we compared both against independent representations of gene function. Gene2vec^52^ generates embeddings from transcriptional data from Gene Expression Omnibus, supplemented with MSigDB^53^ pathway annotations; HiG2vec^54^ derives purely from Gene Ontology hierarchies in hyperbolic space. CTS outperforms co-expression on both, but its advantage is substantially greater with HiG2vec (correlation 0.26 vs. 0.06) than Gene2vec (0.262 vs. 0.220) (**Fig. 3C**). Permutation tests confirm that correlations with CTS are significant (*p* = 0.0001, **Methods**). The comparably stronger agreement with a purely functional representation, HiG2vec, indicates CTS captures gene relationships beyond expression correlation.

The power of CTS is particularly clear in gene families whose members concentrate within a few TADs. In olfactory receptors and protocadherins, CTS substantially exceeds co-expression (olfactory receptors: 0.099 vs −0.002; protocadherins: 0.196 vs 0.001 after median centering; **Fig. 3E-G, Methods**). Histone genes, coordinately regulated during DNA replication^55^, show even higher CTS (0.251 vs 0.021) with modestly better CTS–co-expression agreement (0.114). All differences are highly significant (*p* < 10^−15^; **Fig. 3H**); keratin genes, by contrast, show strong concordance (**Fig. S3**). These patterns suggest TADs may group genes by shared transcriptional context rather than co-expression, a hypothesis we test directly in the next section.

### TADs serve as loci of enriched transcriptional context

Gene pairs sharing a TAD show elevated transcriptional similarity. We defined TAD and non-TAD gene pairs as above and computed the CTS and co-expression within each set of pairs (**Fig. 4A**). Both measures were significantly higher for TAD pairs (*p* < 10^−15^, one-sided t-test), but the effect was markedly stronger for CTS (Cohen’s d = 0.28 vs 0.12), confirming that CTS better captures relationships associated with TAD co-membership.

**Figure 4:**
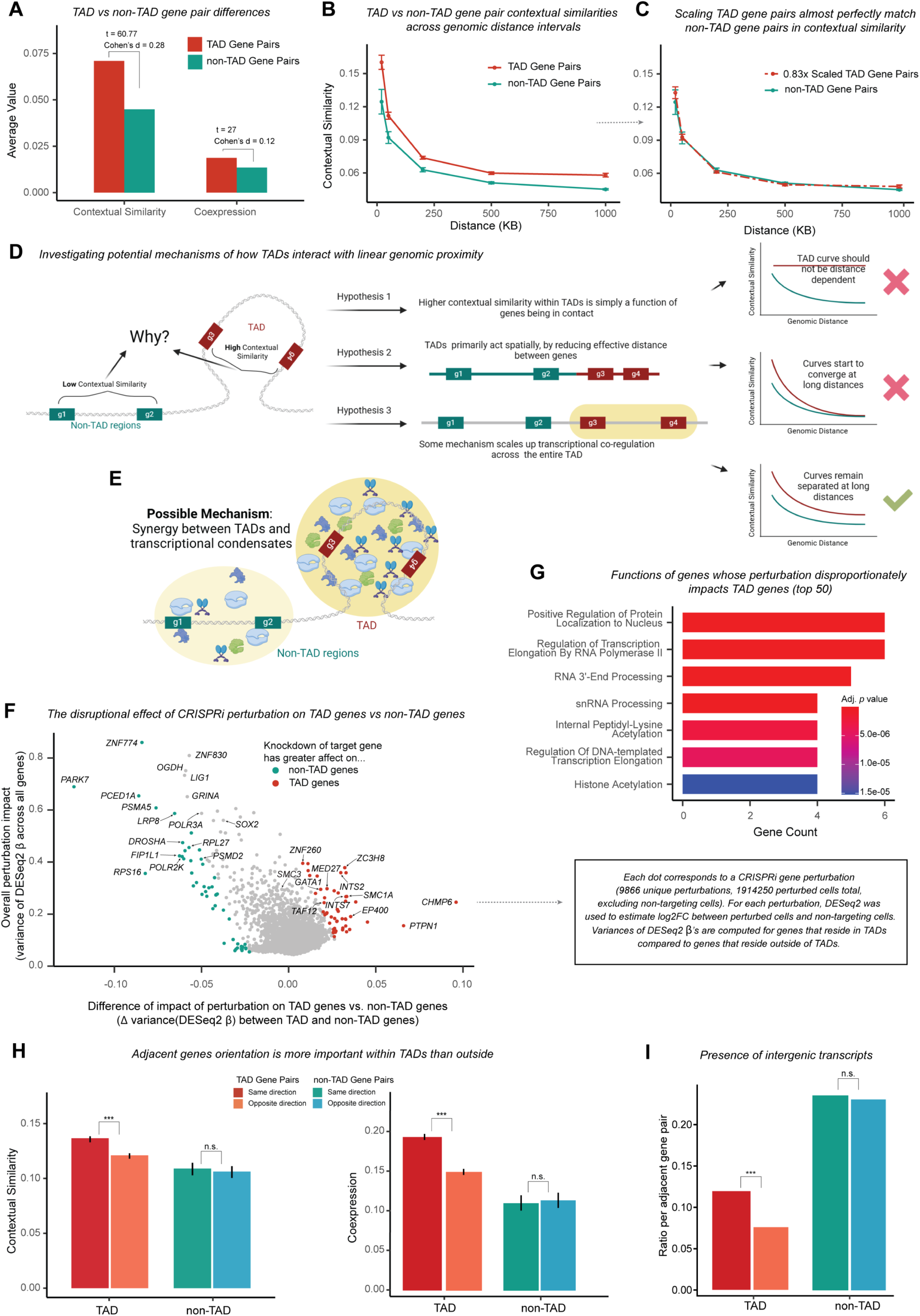
Comparing CTS in TAD and non-TAD gene pairs provides clues into TADs’ role in transcription. **A**) Using the TAD Map, we classified gene pairs on the same chromosome within 2 megabases as TAD or non-TAD pairs based on whether they share a TAD. TAD gene pairs show significantly higher CTS and co-expression than non-TAD pairs, with CTS showing a more pronounced difference. **B, C**) To control for genomic distance effects, we binned gene pairs by distance and compared their average CTS. While TAD gene pairs consistently show higher CTS than non-TAD pairs across all distances (B), both groups show similar distance-dependent decline. Scaling TAD pair CTS by 0.83x matches non-TAD levels (C), revealing that TADs enhance CTS by 20.5% multiplicatively across all genomic distances. **D, E**) We considered three hypotheses for how TADs enhance gene co-regulation: (1) TADs might create distance-independent contacts between all genes, which would result in constant CTS across distances; (2) TADs might reduce effective distances between genes, which would lead to convergence of TAD and non-TAD CTS at long distances; or (3) TADs might work through a mechanism that multiplicatively enhances co-regulation across all distances. Our observations support the third hypothesis, suggesting a TAD-wide regulatory mechanism. Phase-separated transcriptional condensates, which can create specialized compartments spanning large genomic regions, represent a compelling candidate for such a mechanism, leading us to hypothesize that TADs work synergistically with condensates to create specialized regulatory environments. **F, G**) To identify factors regulating TAD-specific transcription, we analyzed a genome-wide Perturb-seq dataset^69^. For each CRISPRi perturbation, we used DESeq2 to estimate log2-fold-changes (*β*) between perturbed and control cells. We then identified perturbations that induced significantly different transcriptional variance in TAD versus non-TAD genes (F). Among the strongest hits were condensate-relevant genes including TAF12, INTS2, and INTS7. Gene Ontology analysis of TAD-specific regulators revealed enrichment for RNA-processing and transcriptional control pathways, consistent with condensate-mediated regulation (G, also see Fig. S3). **H**) A condensate-based mechanism of transcriptional regulation would predict easier co-transcription of same-orientation versus opposite-orientation adjacent genes. Indeed, within TADs, both CTS and co-expression are significantly higher for same-orientation gene pairs (*p* < 10^−11^), while no such difference exists outside TADs. **I**) Similarly, intergenic transcripts, indicators of errorneous transcriptional read-through, are significantly more frequent between same-orientation genes within TADs (*p* < 10^−11^), but show no orientation bias outside TADs.

The TAD effect persists across genomic distance scales, even after accounting for the known tendency of genomically-proximal genes to share functional relationships^56, 57^. We stratified our analysis by distance, grouping gene pairs by their pairwise distance: less than 20 kb, 20–50 kb, 50–200 kb, 200–500 kb, and 500–1000 kb. TAD pairs showed consistently higher CTS across all intervals (*p*-values of 6.04 × 10^−8^, 4.60 × 10^−10^, < 10^−15^, < 10^−15^, and < 10^−15^ respectively) (**Fig. 4B**), with the relative magnitude of this difference preserved even as CTS decreased with distance. To ensure robustness and mitigate scFM-specific bias, we repeated the CTS analysis using another scFM, Transcriptformer^58^, and observed the same pattern (**Fig. S4**). Our findings contrast with Long et al.^59^, who reported no TAD-specific effects beyond proximity using co-expression analysis. Indeed, when we repeated our distance-stratified analysis using co-expression (**Fig. S5**), the TAD-specific signal was substantially weaker, suggesting their conclusion reflected the limitations of the co-expression measure. We further extended this co-expression analysis to 700 additional single-cell transcriptomic datasets from scBaseCount^60^, spanning 18 million cells across 40 tissues, and found similar results (**Fig. S6**).

The structure of CTS’s distance dependence constrains possible physical mechanisms (**Fig. 4D-E**). If TADs created isolated compartments overriding genomic proximity effects, CTS within TADs would be distance-independent. It is not. If TADs acted solely through chromatin compaction (i.e., they uniformly reduce the effective genomic distance), at very long genomic distances TAD and non-TAD CTS would converge. They do not. Instead, we observe a third pattern: TAD membership multiplicatively enhances CTS while preserving distance dependencies. Non-TAD pairs consistently showed 83% of the CTS observed in TAD pairs across all intervals, corresponding to a multiplicative enhancement of 20.5% (**Fig. 4C**). We determined this scaling factor by minimizing the area between CTS curves for TAD and appropriately scaled non-TAD pairs (**Methods**).

What physical process might produce this multiplicative enhancement? One candidate is transcriptional condensates (TCs): dynamic, membrane-less compartments formed by liquid-liquid phase separation that concentrate transcriptional machinery^61–64^. Phase-separated compartments naturally create multiplicative effects on local concentrations^65–68^, consistent with our observations. Individual TCs may be too small to span entire TADs; if this model is correct, either larger “TAD condensates” or the collective action of multiple smaller TCs could contribute to the observed effects. We emphasize that this remains a hypothesis requiring direct experimental validation. Nonetheless, if TADs and condensates do interact systematically, indirect signatures should be detectable in existing genomic data, as we investigated next.

### Systematic regulation in TAD regions differs mechanistically from that of non-TAD regions

If TADs are associated with specialized transcriptional environments, this should manifest in diverse transcriptional readouts. Leveraging publicly available data, we investigated three specific predictions: (1) perturbing regulatory genes should differentially affect transcription within versus outside TADs; (2) gene orientation should constrain co-regulation more strongly within TADs, where shared machinery may be spatially concentrated; and (3) transcriptional read-through between adjacent genes should similarly depend on orientation within TADs.

For the first analysis, we examined genome-wide Perturb-seq data from Replogle et al.^69^, where 9,866 genes were knocked down via CRISPRi with scRNA-seq readout. For each perturbation target, we quantified transcriptional disruption separately for genes inside and outside TADs (**Fig. 4F, Methods**). Extending the analysis to other studies—RPE1, HepG2, K562 essential and Jurkat datasets^70^— yielded broadly consistent patterns across cell types (**Fig. S7, Methods**).

Our analysis framework recapitulates known regulators of transcription and TAD organization. Perturbations causing the most widespread disruption, regardless of genomic location, were enriched for core transcriptional processes such as RNA Polymerase II elongation and initiation (GO:0034243, Adjusted p = 2.39 × 10^−6^; GO:0060261, Adjusted p = 6.09 × 10^−6^)^71^. Mediator complex genes (MED12, MED30, MED21, MED9), which play a central role in transcriptional regulation^72, 73^, ranked in the 99th percentile (**Fig. S8A**)^73, 74^. Perturbations of known TAD-organizing factors showed the expected TAD-specific effects. Cohesin subunits SMC1 and RAD21, which mediate loop extrusion^75–77^, produced strong differential disruption between TAD and non-TAD regions (99th and 97th percentiles, respectively). SMC3 perturbation affected both regions strongly, consistent with its genome-wide role. CTCF perturbation yielded broadly TAD-specific effects (75th percentile), in line with its insulator function at boundaries^2, 36^. These results confirm the framework’s sensitivity to both global transcriptional regulators and TAD-specific organizers.

Among perturbations preferentially affecting TAD regions, we found enrichment for histone acetylation factors (GO:0016573, Adjusted *p* = 4.074 × 10^−4^), including TAF12, ACTL6A, EP400, and DMAP1 (**Fig. 4G**; full list in **Fig. S8B**). These findings align with prior work: loss of H4 acetylation increases long-range contacts beyond TADs^78^, and TAFs contribute to phase-separated transcriptional hubs^79^. A second category, the Integrator complex (INTS2, INTS5, INTS7, INTS8)^80^, also showed TAD-specific enrichment, with roles in snRNA and ncRNA processing (GO:0016180, Adjusted *p* = 1.713 × 10^−5^; GO:0034470, Adjusted *p* = 1.614 × 10^−3^).

Perturbations disproportionately affecting non-TAD regions were enriched for ribosomal processes (**Fig. S8C**), including Ribosome Biogenesis (GO:0042254, Adjusted *p* = 1.254 × 10^−15^) and rRNA Processing (GO:0006364, Adjusted *p* = 3.536 × 10^−6^), with multiple RPS and RPL subunits among the top hits (**Fig. S8D**). We note that non-TAD genes are fewer and already enriched for ribosomal function, which may inflate these estimates. Nonetheless, the pattern suggests a possible division of labor, with nucleoli, membrane-less condensates enriched near low-gene-density heterochromatin^81, 82^, preferentially organizing ribosomal genes outside TADs.

#### Gene orientation constrains transcriptional relationships within TADs

If adjacent genes share transcriptional machinery more readily within TADs, orientation should matter: same-strand pairs would have easier access to shared machinery than oppositely oriented pairs. We tested this using CTS. Within TADs, same-orientation pairs indeed exhibited significantly higher contextual similarity (0.137 vs. 0.121, *p* = 4.5 × 10^−14^); outside TADs, orientation had no effect (0.109 vs. 0.106, *p* = 0.508) (**Fig. 4H, Methods**). Co-expression analysis yielded consistent results: higher correlation for same-orientation pairs within TADs (0.19 vs. 0.15, *p* < 10^−15^) but no difference outside.

In our third analysis, transcriptional read-through—when intergenic regions are erroneously transcribed—shows the same pattern. Analyzing 11,417 intergenic transcripts from Agostini et al.^83^ (**Methods**), we found that within TADs, same-orientation gene pairs showed significantly more read-through than opposite-orientation pairs (11.9% vs 7.6%, *p* < 10^−15^, one-sided binomial test; **Fig. 4I**). Outside TADs, orientation had no effect, though overall read-through frequencies were higher, suggesting elevated transcriptional noise. These orientation-dependent patterns within TADs, but not outside, are consistent with models where TADs facilitate shared access to transcriptional machinery.

### The TAD Map and CTS framework enables new aging and cancer-related investigations

The TAD Map framework enables systematic investigation of chromatin’s role in cellular state transitions without requiring additional Hi-C experiments for each cell type^84, 85^. Prioritized hypotheses can then be experimentally tested.

#### Cell differentiation

A central question in developmental biology is how cells modulate gene expression while balancing stability and plasticity. Recent work by Pollex et al.^86^ shows that developmental shifts from permissive early enhancer–promoter contacts to more constrained later topologies likely reflect corresponding changes in TAD-level transcriptional organization^12^. We analyzed diverse differentiation trajectories and observed a clear relationship between developmental stage and TAD-mediated gene organization. Using developmental scRNA-seq transcriptomes curated by Gulati et al.^87^, we found that cells in earlier differentiation stages showed significantly stronger expression clustering within TADs (21 of 24 trajectories, **Fig. 5A**, *p* = 0.00019, one-sided t-test) and designed a bootstrap-based “TAD usage” score to quantify this effect (**Fig. S9A-B, Methods**). This pattern persisted across stage definitions and tissues (**Fig. S9C**), suggesting that TAD organization is especially prominent during developmental plasticity.

**Figure 5:**
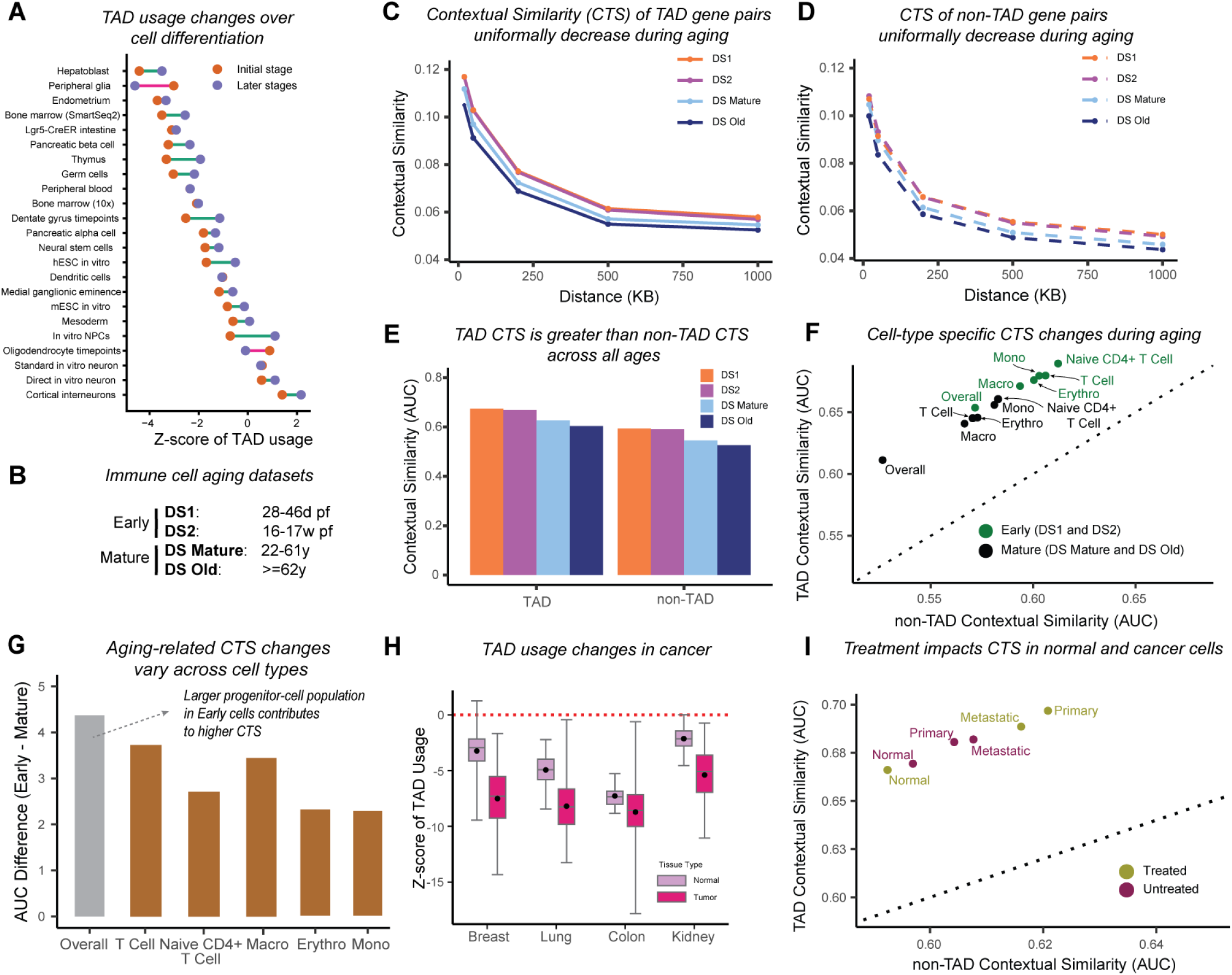
TAD-mediated transcriptional organization changes systematically during development, aging, and cancer. **A**) Analysis of 24 non-branching cell differentiation trajectories^87^ reveals that early-stage cells consistently show higher TAD usage (more TADs with non-zero expression activity), indicating stronger transcriptional clustering (also see Fig. S4). **B**) To study aging effects, we analyzed immune cells from two large datasets^91, 92^, spanning prenatal (DS1, DS2), mature adult (DS Mature), and older adult (DS Old) stages. **C, D, E**) Fine-tuning scGPT on each age group and recalculating CTS revealed systematic changes with age. Both TAD (C) and non-TAD (D) gene pairs show decreasing CTS with age across all genomic distances. Area under the curve (AUC) analysis confirms this decline while showing that TAD pairs maintain higher CTS than non-TAD pairs across all age groups (E). **F, G**) Cell-type specific analysis reveals that aging-related chromatin changes occur not just through shifts in cell populations but also within specific cell types (particularly T cells and macrophages). Combining early (DS1, DS2) and mature (DS Mature, DS Old) datasets, we found that while early cells generally show higher CTS (F), the magnitude of age-related decline varies substantially across cell types (G), with T cells and macrophages showing the strongest changes. **H**) Analysis of The Cancer Genome Atlas (TCGA) data shows that, like early developmental stages, tumor cells exhibit increased TAD-based expression clustering across multiple cancer types. **I**) In colorectal cancer, 5-fluorouracil chemotherapy induces opposing changes in chromatin organization between normal and cancer cells: treated tumor cells show increased CTS while treated normal cells show decreased CTS, suggesting distinct adaptive responses to treatment stress.

#### Aging

Motivated by this developmental pattern, we asked whether TAD-level organization also shifts with age. We used the immune system as a model, where aging alters lineage distributions and chromatin states^88–90^, to test whether these changes reflect solely shifts in cell type proportions or also intrinsic within–cell-type changes in chromatin regulation. We analyzed scRNA-seq datasets from human immune cells spanning prenatal to late adult stages^91, 92^ and grouped samples into four age categories (DS1: 28–46 days post-fertilization; DS2: 16–17 weeks post-fertilization; DSmature: 22–61 years; DSold: over 61 years) (**Fig. 5B**) to assess whether CTS between gene pairs changes systematically with age.

As cells age, we observed a decline in the proximal transcriptional programs and machinery coordinating gene pairs using CTS area-under-curve metric. For each curated dataset (DS1, DS2, DSmature, DSold), we fine-tuned the whole-human scGPT model, extracted updated gene embeddings, and computed CTS for TAD and non-TAD gene pairs as before (**Fig. 5C–D, Methods**). CTS was highest in DS1, followed by DS2, DSmature, and DSold, a pattern consistent for both TAD and non-TAD gene pairs (**Fig. 5E**). The decline was substantial. In TAD pairs, DS1’s CTS (measured by AUC) exceeded DSold by 11.75%, DSmature by 7.66%, and DS2 by 0.90%. Non-TAD pairs showed similar declines (DS1 exceeding DSold by 12.71%, DSmature by 8.80%, and DS2 by 0.51%). This systematic decline suggests that aging progressively modulates the spatial constraints on transcriptional machinery.

To distinguish between population-level and cell-intrinsic changes, we fine-tuned the model separately on subsets of scRNA-seq data and examined CTS changes in T cells (general and naive thymus-derived CD4-positive, alpha-beta T cell), macrophages, monocytes, and erythrocytes across early and mature stages (**Methods**). We found that CTS decreased with age within each cell type (**Fig. 5F**), with general T cells and macrophages showing the most pronounced decline (**Fig. 5G**). Our cell-type–specific analysis suggests that the age-related decline in CTS arises from both shifts in cell type proportions and intrinsic within–cell-type changes, with the strongest effects in T cells and macrophages, highlighting these populations as key targets for studying how aging modulates chromatin organization.

#### Cancer

If TAD-mediated transcriptional clustering covaries with developmental plasticity, cancer cells should also exhibit the elevated TAD usage typical of progenitor-like states, as they often dedifferentiate or fail to fully differentiate. Indeed, analysis of bulk RNA-seq data from The Cancer Genome Atlas (TCGA) across blood, brain, lung, and renal cancers revealed that tumor cells exhibit greater clustering of transcriptionally active genes within TADs compared to normal tissues^93^ (**Fig. 5H**). This difference in TAD usage patterns is robust to adjustments for the higher number of expressed genes in tumor cells (**Methods**).

We also examined how cancer treatment might modulate cell plasticity during disease progression by applying CTS analysis to a colorectal cancer dataset that tracks plasticity through metastasis^94^. After fine-tuning scGPT on this data, we found that in untreated samples, cancer cells (both primary tumor and metatstatic) show higher CTS than normal cells, consistent with the TCGA analysis above. However, After 5-fluorouracil–based chemotherapy, tumor cells (primary and metastatic) showed increased CTS relative to untreated tumors, whereas normal cells exhibited a pronounced CTS decrease (**Fig. 5I**).

The divergence of post-treatment CTS patterns between normal and cancer cells align with well-established biological responses to chemotherapy. It indiscriminately targets rapidly dividing cells. In normal tissues, the slightly lower CTS after treatment is consistent with a reduction in the proportion of progenitor cells. In tumors, chemotherapy can select for cancer stem cells (CSCs) that possess inherent resistance mechanisms, including enhanced DNA repair capabilities, drug efflux pumps, and anti-apoptotic pathways^95, 96^. The elevated CTS in treated cancer populations could reflect this enrichment of stem-like cells, which maintain plastic transcriptional programs even under treatment pressure. Although CTS–plasticity correlations fit known biology, causal links between TAD organization and treatment response will require further investigation. If validated, our findings would address a major gap in clinical diagnostics, with a quantitative estimator of post-treatment CSC enrichment.

### TAD signatures highlight heterogeneity of scRNA-seq data

While the unprecedented detail and volume of scRNA-seq data have had a transformative impact on biological research, challenges such as data sparsity, uneven gene coverage (“dropouts”), and platform effects remain^97^. To complement existing dimensionality reduction techniques, we introduce TAD signatures, an epigenetically-informed scRNA-seq representation that leverages TAD Map gene groupings to mitigate technical noise while preserving biological signal, and that can be inferred for any human or mouse scRNA-seq dataset without requiring Hi-C data (**Methods**). Across 193 cancer cell lines, drug response profiles (4,686 compounds from the PRISM screen) correlate with TAD signature–based measures of higher-order transcriptional heterogeneity, indicating that TAD signatures capture phenotypically relevant variability beyond what is accessible through gene-level expression alone (**Fig. S10A**). In addition, TAD signatures improve automated cell type inference by enhancing clustering accuracy (**Fig. S10B**) and enabling reliable discrimination of closely related cell subtypes with subtle transcriptional differences that challenge gene-based approaches (**Fig. S10C**).

## Discussion

This work seeks to reframe how chromatin structure relates to gene expression. We characterize TADs not as deterministic regulators where membership dictates co-expression but as probabilistic biases: genomic contexts where transcriptional coordination is elevated. Any individual TAD may deviate from this pattern, but across the genome—and at all distance scales—genes sharing a TAD show ∼20% higher contextual similarity than genes at equivalent distances outside TADs. The gradual modulation we observe during aging and in cancer is consistent with this probabilistic view: chromatin organization biases transcription without absolutely constraining it.

Two conceptual advances enable this perspective. The TAD Map abstracts chromatin structure to a species-level “bag of genes” representation, tolerating boundary variation while enabling genome-wide analysis without cell type–specific Hi-C. Contextual transcriptional similarity (CTS) leverages single-cell foundation models to capture gene relationships that co-expression misses—particularly functional substitutability among genes that rarely co-express.

CTS exposes a class of gene relationships invisible to traditional analysis. Long et al., limited to co-expression analysis in only two cell types^59^, could not disentangle TAD-specific effects from genomic proximity. Leveraging single-cell foundation models (scFMs), CTS reveals systematic TAD-mediated enhancement that holds across distance scales and cell types. More broadly, scFMs excel at capturing broad-but-subtle biological patterns when combined with prior knowledge of genomic organization. This is particularly powerful in small-data settings. While individual age-stratified datasets were too small for robust correlation estimation, fine-tuning scGPT on each revealed clear distinctions. Our work demonstrates that scFM representations can be repurposed beyond their original framing. While the original scGPT study interpreted gene embedding similarity as simply recapitulating gene regulatory networks^45^, our CTS-based view reveals new patterns the network interpretation misses.

Our observations are consistent with a physical model in which TADs interact with condensates, phase-separated compartments that concentrate transcriptional machinery. Three independent lines of evidence support this possibility: perturbation of condensate-associated genes disproportionately affects transcription within TADs; same-orientation gene pairs show enhanced contextual similarity specifically within TADs; and transcriptional read-through between adjacent genes follows the same orientation-dependent pattern within TADs but not outside them. We note these patterns are correlational. Establishing causality (and its direction) will require direct experimental manipulation, e.g., perturbing condensate formation within defined TADs or imaging studies to localize phase separation in vivo^98, 99^.

Our framework opens new avenues for studying cellular transitions. If TAD-associated organization reflects regulatory coordination, it should track with cellular plasticity. It does. Cells in early differentiation show stronger TAD-level coordination; this organization declines systematically with aging, even within specific cell types like macrophages. Cancer cells show elevated TAD usage, with divergent responses to treatment between tumor and normal populations. Establishing causality will require direct experimental manipulation, but the key advantage of our approach is hypothesis-generation: it reveals these large-scale patterns across diverse conditions without requiring Hi-C data for each cellular state.

Our approach has limitations that suggest future directions. There remains active debate regarding the conservation of TADs across tissues and cell types^24^. While the bag-of-genes model mitigates boundary variability, it does not resolve this broader controversy; our findings should be interpreted within this context. The TAD Map enables species-wide analysis but sacrifices resolution—it cannot capture fine-grained structural variations between cell types, subtle boundary shifts, or chromatin sub-structure such as sub-TADs. Targeted functional studies could probe how specific genomic contexts or cell states deviate from the general patterns we report. Our focus on protein-coding genes leaves open questions about non-coding RNA regulation within TADs. Our treatment of boundary-spanning genes (currently assigned to both flanking TADs) is conservative but deserves dedicated investigation; these genes may experience distinct regulatory environments. Our systematic framework can serve as the base for more refined models of chromatin–transcription synergy. Computational methods for inferring cell-specific 3D genome struc-ture^90, 100–106^ and multimodal sequencing approaches^107–112^ could integrate with scFM representations fine-tuned on individual cell types, allowing hierarchical chromatin organization to inform CTS while accommodating structural variation. Such a hierarchical scheme may also help identify alternative physical mechanisms, such as the nested enhancer network described by Lin et al.^113^. Other epigenetic modalities, such as histone modifications, dynamically interact with chromatin topology^114, 115^ and could further refine the framework.

By bridging genome organization with foundation model-based representation learning, this work enables a systematic investigation of chromatin’s role in cellular transitions without requiring structural data for each state. The TAD Map and CTS framework demonstrates how integrating prior knowledge with foundation models can reveal patterns obscured by cell type–specific analyses. Lastly, our work advocates for reimagining transcriptional regulation as a convolution of probabilistic mechanisms—chromatin 3D structure, chromatin accessibility, transcriptional condensates—that form the background against which discrete actors like transcription factors perform.

## Methods

### Cell type-specific TAD definitions

We sourced cell type-specific TAD architectures from Liu et al.’s TADKB database^33^, selecting TAD definitions inferred with the Directionality Index (DI) technique at 50 kb resolution. We chose TADKB because it ensured a consistent “Hi-C to TAD” mapping across experimental data from multiple studies. We retrieved data for all the human and mouse cell lines available in the database: seven for human (GM12878, HMEC, NHEK, IMR90, KBM7, K562, and HUVEC) and four for mouse (CH12-LX, ES, NPC, and CN). These TAD definitions were computed by Liu et al. using source Hi-C data from Rao et al.^2^ (Gene Expression Omnibus, GEO, accession GSE63525) and Bonev et al.^116^ (GEO accession GSE96107).

### Expanded Cell Type Coverage

We expanded our analysis 2.5-fold by increasing coverage from 11 to 27 cell types, resulting in 17 human and 10 mouse cell types drawn from diverse tissues, developmental stages, and both in vivo and in vitro systems. These additional datasets were sourced from the 4D Nucleome project^34^. For human, the updated compendium of cell types includes GM12878 (B lymphoblastoid), K562 (erythroleukemia), KBM7 (haploid leukemia), Hap1 (haploid myeloid), bone-marrow lymphocytes (primary lymphocytes), HMEC (mammary epithelial), NHEK (epidermal keratinocyte), IMR90 (fetal lung fibroblast), HFF-hTERT (immortalized foreskin fibroblast), HUVEC (umbilical vein endothelial), HepG2 (hepatocellular carcinoma), CyT49-derived gut-tube endoderm, H1-derived pancreatic progenitor, olfactory receptor cells, left-atrium cardiomyocytes, HeLa (cervical adenocarcinoma), and H9-derived cardiomyocytes. For mouse, the expanded set comprises CH12-LX (B-lymphocyte lymphoma), ES (embryonic stem cell), NPC (neural progenitor cell), CN (cortical neuron), cardiac muscle, Sertoli (testicular support) cells, trophoblast (placental lineage), granulosa (ovarian somatic) cells, double-negative alpha-beta thymocytes (immature T-cell precursors), and olfactory receptor cells.

### Reconstruction of TAD Maps

We regenerated both human and mouse TAD Maps using these expanded panels of cell types. For all newly added cell types, TADs were called independently using the Directionality Index (DI) method^26^ and the Insulation Score (IS) method^35^. The resulting architectures were used as inputs to construct revised species-specific TAD Maps. Broadly, these new maps remain in close agreement with our originally reported TAD Maps, indicating that increased cellular diversity and variation in TAD-calling methodology do not materially alter the large-scale TAD architecture.

### Reference genome versions

All genomic coordinates and gene names correspond to Ensembl v102, with the human and mouse reference genomes being hg38 and mm10, respectively. With the *liftOver* program^117^, TAD definitions for human cell types were mapped to the hg38 reference genome from hg19, the reference genome used in TADKB. TAD definitions for mouse cell types already corresponded to the mm10 reference genome.

### Maximum likelihood estimation of the consensus TAD scaffold

We infer TADs independently for each chromosome: our algorithm takes as input a list of cell type-specific TAD architectures for the chromosome and outputs the consensus TAD definition. Both the input and output definitions specify TADs as a set of non-overlapping genomic intervals along the chromosomes. Our algorithm currently operates at a 50 kb resolution with both the input and output defined at that granularity.

We divide the entire chromosome into segments of size R (currently, R = 50 kb) and for every possible pairwise combination of segments, compute the likelihood that the two segments share a TAD. More formally, given segments at loci *i* and *j*, we define *X*_*i* *j*_ as the number of input cell type-specific TAD architectures in which *i* and *j* share a TAD. Our goal is to infer *C*_*i* *j*_, where *C*_*i* *j*_ ∈ {0, 1} indicates if *i* and *j* share a consensus TAD. Additionally, *C*_*i* *j*_ need to obey integrity constraints that correspond to a valid TAD architecture; we describe these later. To accommodate the differing amounts of input data available for human and mouse in a unified framework, we discretized *X*_*i* *j*_ into three levels: *X*_*i* *j*_ ≥ *T*, *T* > *X*_*i* *j*_ > 0, and *X*_*i* *j*_ = 0. With data for 7 human cell types and 4 mouse cell types, we chose *T* as 4 (for human) and 2 (for mouse), so that the three levels of *X*_*i* *j*_ express the intuition that it receives support from the majority, at least one, or none of the input cell types, respectively. We then parameterize our likelihood model *P*(*X*_*i* *j*_ |*C*_*i* *j*_) as follows:

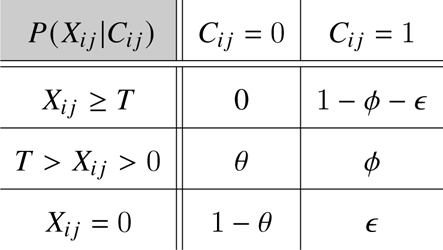

where 0 < *θ*, *ϕ*, *ε*, (1 − *ϕ* − *ε*) < 1. Our parameterization implies that a segment pairs *i*, *j* which share a TAD in a majority of the input cell types (*X*_*i* *j*_ ≥ *T*) must be present in a consensus TAD. On the other hand, we provide for the possibility (*ε* > 0) that a segment pair *i*, *j* with no support from any of the input architectures might still share a consensus TAD— this allows us to stitch together overlapping TAD ranges across cell type-specific inputs if strong overall support exists for a broad TAD at that locus.

Under our model, the likelihood of the observations **X** = {*X*_*i* *j*_ } given the consensus TAD scaffold **C** = {*C*_*i* *j*_ } is

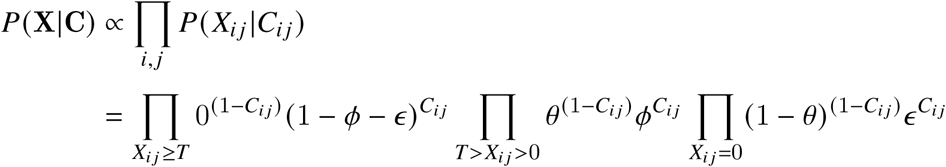

The first sub-product imposes the binding constraint that *C*_*i* *j*_ = 1 for all segment pairs {*i*, *j* | *X*_*i* *j*_ ≥ *T*}. Given that, we can focus on the remaining two sub-products to maximize the log-likelihood:

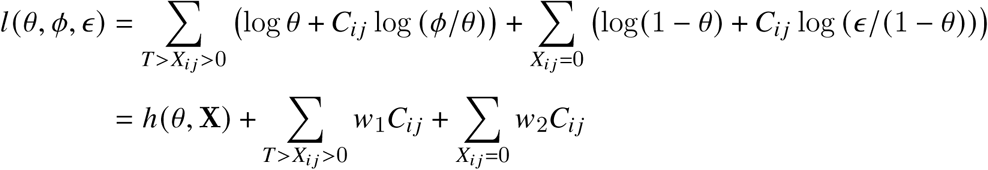

where ℎ(*θ*, **X**) is not a function of *C*_*i* *j*_, and the terms *w*_1_ and *w*_2_ represent more convenient combinations of the parameters *θ*, *ϕ*, and *ε*. Maximizing the log-likelihood thus requires solving

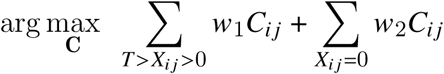

We note that integrity constraints on *C*_*i* *j*_ link the terms: **C** needs to be transitive, i.e., if segments pairs (*i*, *j*) and (*j*, *k*) share a TAD then so must (*i*, *k*). Also, a TAD must be contiguous: if *C*_*i* *j*_ = 1 then *C*_*ij*_ = *C*_*j* *j*_ = 1 for all segments *v* between *i* and *j*. Intuitively, this formulation describes a trade-off between biasing towards long TADs, which cover more true positive *C*_*i* *j*_ s (guided by *T* > *X*_*i* *j*_ > 0 cases) but also have more false positives (driven by *X*_*i* *j*_ = 0 cases), and short TADs where the false positives will be fewer but the risk of false negatives increases.

To maximize *l* and compute **C**, we formulated a dynamic programming algorithm that splits the chromosome into recursively smaller ranges and finds the globally optimal combination of *C*_*i* *j*_ assignments. We chose *w*_1_ = 0.5, *w*_2_ = −1 for human and *w*_1_ = 0.05, *w*_2_ = −3 for mouse; these choices produced consensus TAD scaffolds where the number of TADs and the distribution of their lengths was in line with the corresponding statistics for cell type-specific TAD architectures.

### TAD boundaries

As an alternative quantification, we compare these estimates to a null hypothesis where TAD boundaries are independently dispersed between gene boundaries in each cell type while preserving the number of TADs. As (**Fig. 2C**) shows, for a gene pair that co-occupies a TAD in one of the architectures, the expected number of other cell-type architectures that agree with this grouping under the null hypothesis is just 0.85 (human) and 0.29 (mouse), implying that the extent of agreement actually observed is highly significant (one-sided binomial test, *p* = 7.3 × 10^−75^ for human, 1.7 × 10^−4^ for mouse).

### Agreement with CTCF ChIP-seq

We sourced CTCF ChIP-seq data from the ENCODE database^39^, consisting of 281 human and 28 mouse studies (comprising 665 and 109 replicates, respectively). These studies covered many cell types where concordance of TADs and CTCF binding sites was previously uncharacterized because Hi-C data for the cell type was unavailable. We filtered to only keep peaks with Irreproducible Discovery Rate (IDR) less than 0.05 and mapped these peaks to our estimated TAD scaffold. We then grouped the peaks by their location relative to TADs, partitioning the entire genome into the following disjoint categories: *TAD Boundary* (50 kb segments on each side of the TAD), *Inside Boundary* (two 50 kb segments inside the TAD, just interior to the TAD boundaries), *Outside Boundary* (two 50 kb regions outside the TAD, just exterior to the TAD boundaries), *TAD Interior* (the part of the TAD that’s not in the *Inside Boundary* segments), and *TAD Exterior* (all other parts of the genome). With the CTCF peak widths being much smaller (median width = 273 bp) than the granularity of our TAD scaffold (50 kb resolution), we assumed that a peak would not span two segments and assigned each peak to its genomic segment based only on the peak’s midpoint locus. Finally, we counted the number of peaks in each segment category, normalizing that count by the aggregate length of genomic segments in that category.

Using the ChIP-seq data, we also assess the precision of our inferred TAD boundaries. We evaluate binding prevalence in 50 kb regions on either side of our predicted TAD boundary (we recall that our TAD scaffold is inferred at a 50 kb resolution). Supporting our inference, we find that the regions adjoining the boundary have substantially lower CTCF binding rates than at the TAD boundary itself. However, the CTCF binding rate in these adjoining regions is still higher than the rate observed in the TAD exterior or interior. This could be explained by minor boundary variations across cell types (as is indeed known to happen) or if the “effective” TAD boundary is wider than our 50 kb definition. We note that the bag-of-genes model for TADs is designed to handle such ambiguity: as long as the gene memberships in a TAD are unchanged, minor variations in its boundaries are immaterial.

### Expression quantitative trait loci

From the GTEx Analysis Release V8^40^, we sourced data for all 49 tissues with available single-tissue eQTL data. We filtered the data, limiting ourselves to eQTLs with p-value less than 10^−5^. For each gene and eQTL pair, we computed the genomic distance between the transcription start site (TSS) of the gene and the eQTL locus; based on it, we assigned the ⟨ TSS, eQTL ⟩ pair to one of the genomic-distance bins partitioned by the following cut-points (all units in kb): [0, 5, 10, 20, 30, 50, 75, 100, 150, 250, 350, 450, 550, 650, 750, 1000]. We limited the upper bound of our evaluation range to 1 Mb, since the GTEx corpus limits single-tissue eQTL reports to this range. We counted the number of observed pairs in each bin, separately tracking pairs where the TSS and eQTL loci shared a TAD and pairs where they did not; we additionally separated pairs where the eQTL locus was upstream of TSS from those where it was downstream. The probability *p*(*d*) of an eQTL occurring at a distance *d* from the TSS was then estimated by dividing the number of observed ⟨TSS, eQTL⟩ pairs in each bin by the midpoint of the bin’s genomic range.

### Comparison of contextual transcriptional similarity (CTS) with co-expression

We com-puted CTS by leveraging the whole-human scGPT model, accessed from its GitHub reposi-tory (https://github.com/bowang-lab/scGPT). Gene embeddings were extracted from the “encoder.embedding.weight” matrix, using gene row mappings provided in the accompanying vocab JSON file. Chromosome and gene location data were obtained from the hg38 reference genome. For all gene pairs located on the same chromosome and within 2 megabases of genomic distance, we calculated Pearson correlations between their embedding vectors, defining these correlations as the CTS for each pair.

To calculate co-expression, we analyzed cell atlas datasets downloaded from CELLxGENE (https://cellxgene.cziscience.com/datasets), focusing on the Human Breast Cell Atlas, Human Brain Cell Atlas, Human Cell Atlas of Fetal Gene Expression, and Human Heart Atlas. Raw transcript counts for each cell were normalized to a total of 10,000 and subsequently log-transformed using log1p. For genes present across all datasets, we computed Pearson correlations of expression values for gene pairs on the same chromosome within 2 megabases. Finally, we compared CTS and co-expression, analyzing how these measures differ in capturing gene relationships.

### Gene family analysis

To investigate gene family-specific patterns, we filtered gene pairs such that both genes belonged to one of the families of interest: olfactory receptors, protocadherins, histones, or keratins. After calculating CTS and co-expression, we highlighted gene pairs from specific families and compared their values to the broader distribution (**Fig. 3E-H**).

We also found genes in certain gene families primarily reside in the same few TADs. Olfactory receptors are plentiful in TADs on chr11-4150000-5450000, chr11-55000000-56850000, and chr11-123750000-124600000. Protocadherins primarily reside in TADs on chr5-140750000-141600000. Histones primarily reside in TADs on chr6-26000000-26600000 and chr6-27150000-27950000. Lastly, keratins primarily reside in TADs on chr17-40600000-41650000, chr21-30300000-31100000 and chr12-52150000-53050000.

### Comparison of CTS to other gene function embedding similarity scores

We compared CTS to other gene function embedding similarity scores to determine whether scGPT gene embeddings capture functional information beyond gene expression. We included Gene2vec, which generates 200-dimensional vectors using co-expression patterns and functional gene sets, and HiG2Vec, which creates 1000-dimensional Poincaré embeddings to represent the hierarchical structure of Gene Ontology and gene semantics. To assess whether the association between HiG2Vec and CTS could be explained by chance, we performed a permutation test in which CTS values were shuffled 10,000 times and correlations were recalculated. The p value was then computed using the observed correlation. We obtained Gene2vec embeddings from https://github.com/jingcheng-du/Gene2vec and HiG2Vec embeddings from https://github.com/JaesikKim/HiG2Vec. For both methods, we computed gene pair similarities using Pearson correlation and related these similarities to CTS, focusing on gene pairs common to both subsets.

### CTS and co-expression for TAD vs non-TAD gene pairs and across genomic distance intervals

We categorized gene pairs as TAD pairs if they resided in the same TAD based on the TAD Map; otherwise, we classified them as non-TAD pairs. To determine genomic distance, we averaged the distances between the start and end positions of the two genes. We stratified all gene pairs into genomic distance intervals of 0–20 kb, 20–50 kb, 50–200 kb, 200–500 kb, and 500–1000 kb, with intervals defined as up to but not including the upper bound. We grouped gene pairs by TAD status and interval category and calculated their average CTS and co-expression.

### Determining the scaling factor that aligns TAD gene pairs with non-TAD gene pairs

We examined how uniformly scaling CTS for TAD gene pairs best aligns with CTS for non-TAD gene pairs. Across all genomic distance intervals, CTS for TAD gene pairs consistently exceeded that for non-TAD gene pairs. We connected average CTS values per interval to form curves and calculated the area under the curve (AUC) using trapezoidal integration. To evaluate the impact of TADs, we scaled the average CTS for TAD gene pairs iteratively by factors ranging from 0 to 1 while keeping CTS for non-TAD gene pairs fixed. For each scaled TAD CTS, we computed the difference between its AUC and the fixed non-TAD AUC. The scaling factor that minimized the difference in AUCs quantified the effect of TADs on gene pair CTS.

### Perturb-seq to identify targets impacting gene expression within and outside TADs

We analyzed the genome-wide Perturb-seq dataset (https://gwps.wi.mit.edu/) from Replogle et al.^69^, generated using the K562 chronic myeloid leukemia cell line. We used this dataset to identify genetic perturbations that most disrupt overall transcription, as well as transcription specifically in genes located within TADs and those outside TADs. The dataset has 9866 unique CRISPRi gene-targeted perturbations and 1,914,250 total gene-targeted perturbed cells, with 75,328 non-targeting cells. For each perturbation, we used DESeq2^118, 119^ to calculate fold changes in all captured gene expression relative to a randomly downsampled group of non-targeting cells. We used TAD Map and Ensembl v102 gene annotations to categorize all protein-coding genes as either TAD genes (those residing in a TAD) or non-TAD genes (those outside TADs). Transcriptional disruption was quantified as the variance of DESeq2 log2 fold-change (*β*) values for protein-coding genes, calculated separately for TAD and non-TAD genes. We repeated this analysis five times with different randomly downsampled non-targeting groups and averaged the variance metrics to evaluate the perturbation-specific transcriptional impacts on all genes, TAD genes and non-TAD genes.

### Expanded Perturb-seq analysis across multiple cell lines

To broaden our analysis of perturbation-induced transcriptional effects, we incorporated additional Perturb-seq datasets from HepG2 and Jurkat cell lines^70^, as well as from K562 and RPE1 cell lines^69^. We restricted all analyses to the subset of perturbations and genes that were shared across the four datasets, allowing a unified comparison across cell lines. Using the same analytical framework described above, we quantified, for each perturbation and in each cell line, the relative disruption of transcription in genes residing within TADs versus those outside TADs. We then identified perturbations that showed consistent TAD-specific or non–TAD-specific transcriptional effects and focused on those that recurred in at least three of the four cell lines.

### CTS and co-expression in adjacent genes

We use gene information from Ensembl v102 to denote if gene pairs are adjacent or neighoring gene pairs, as well as if they reside on the same strand or if they reside on opposite strands. For analysis with CTS, we used the same gene pairs from previous analysis. For our analysis with adjacent gene pairs For co-expression, we tested on 794 (human) and 69 (mouse) bulk RNA-seq datasets sourced from the ENCODE database (again encompassing a variety of cell types).

### Intergenic transcripts

We sourced data from Agostini et al.’s study of intergenic transcripts. They collected and analyzed data from 38 publicly available datasets, covering over 2.5 billion uniquely mapped reads. We limited ourselves to 11,417 intergenic transcripts that are currently unannotated (the others corresponded most frequently to non-coding RNA fragments). As in the original study, each transcript was mapped to its adjoining genes as per the Gencode 27 reference. We further annotated each intergenic transcript by whether its adjoining genes were a) shared a TAD, and b) if they were oriented similarly.

### Bootstrap test for clustering of expressed genes into TADs

The bootstrap test operates on data from a single cell (in scRNA-seq data) or a single tissue (in bulk RNA-seq data). Given scRNA-seq readout from any cell, we compute *k*, the number of genes with non-zero transcript counts in the cell. Treating gene activity as a binary event, we then generate 500 bootstrap samples of single-cell gene expression in each of which we randomly choose *k* protein-coding genes to be active. For both the actual observation and the bootstrap samples, we map these genes to the TAD Map, computing the number of TADs *n*(*p*, *k*) which have *p* or more active genes; here, *p* = 1 corresponds to identifying the set of TADs with non-zero usage. We estimate the mean and standard deviation of the distribution *n*(*p*, *k*) from the bootstrap samples and, using that, compute the z-score for the actual observation. In bulk RNA-seq data, which includes gene expressed aggregate from a collection of cells, almost all genes have some non-zero expression. There, we pre-set a threshold *k* (say, 5000) and limit ourselves to the top-*k* genes by transcript count; the bootstrap test and z-score are computed for this *k*.

In the test above, if a gene spans two TADs we count it in both TADs. This avoids us having to assign the gene to one TAD or the other arbitrarily. We also believe this to be the conservative choice for our clustering test: it will lead to more TADs per gene— i.e., lower clustering of expressed genes into TADs— than if we were to assign the gene to just one TAD or the other.

### scRNA-seq cell differentiation datasets

We sourced scRNA-seq data from the CytoTrace database made available by Gulati et al.^87^. The study collected and curated scRNA-seq cell differentiation studies across multiple species, covering a variety of protocols and tissues. We chose this corpus as our scRNA-seq testbed, since it covers a diversity of protocols and tissues and allowed us to extend our analysis of also study TAD usage during cell differentiation. Of the 43 datasets available on cytotrace.stanford.edu (while their webpage lists 47 entries, 4 rows are blank), we filtered out studies with fewer than 200 cells and those that did not originate from human or mouse tissue, leaving us with 33 studies. We had difficulty converting 3 of these from the original *RDS* format to a *Scanpy*-compatible format and limited ourselves to the remaining 30 (which covered 70,243 cells); these formed our scRNA-seq corpus.

### Designation of early vs. late-stage cells during differentiation

When analyzing TAD usage during cell differentiation, we further limited ourselves to the 24 scRNA-seq datasets where the putative differentiation trajectory did not have any branches, allowing us to reliably order cells along a differentiation time course. We used Gulati et al.’s annotations of differentiation stage in each study and considered two measures of early versus late differentiation: 1) consider only the cells at the first differentiation stage (*order* = 0 in CytoTrace) as “early”, with all other cells comprising the “late” stage, or 2) partition the cells in each dataset by *order* so that cells are divided roughly equally between the first (“early”) and second (“late”) halves.

### CTS for immune cell aging datasets

We analyzed immune cell scRNA-seq data from Suo et al.^91^ and the single-cell transcriptomic atlas^92^, both sourced from CELLxGENE. From Suo et al.’s dataset, we selected cells whose development stage is in Carnegie stages 13, 19, and 20, designating this subset as DS1. Cells from the 16th and 17th weeks post-fertilization development stage were filtered and grouped as DS2. For the mature immune cell atlas, we included cells from donors aged 22 to 61 years, labeled as DS Mature, and those from donors older than 62 years, labeled as DS Old. To ensure consistency across datasets, we downsampled each to 50,000 cells using Geosketch^120^.

For each dataset (DS1, DS2, DS Mature, DS Old), we fine-tuned pre-trained whole-human scGPT model on a masked gene expression prediction task for 10 epochs. During fine-tuning, all scGPT parameters were frozen except for the gene embeddings (“encoder.embedding.weight”). We adopted default hyperparameters provided in the scGPT tutorials (https://github.com/bowang-lab/scGPT/tree/main/tutorials). Following fine-tuning, we extracted updated gene embeddings and calculated contextual transcriptional similarity (CTS) for the same gene pairs analyzed earlier. To compare overall CTS across datasets, we evaluated CTS trends across genomic distance intervals (**Fig. 5C-D**) and computed the area under the curve (AUC) for TAD and non-TAD gene pairs (**Fig. 5E**) using trapezoidal integration.

### CTS for cell-type specific immune cell datasets

We combined DS1 and DS2 datasets to define the “Early” developmental stage cells and combined DS Mature and DS Old datasets to define the “Mature” stage cells. Within these aggregated stages, we identified five cell types containing more than 2,500 cells: general T cells, naive thymus-derived CD4-positive, alpha-beta T cells, macrophages, monocytes, and erythrocytes. For each cell type in both Early and Mature stages, we fine-tuned the scGPT model for 10 epochs with masked gene expression prediction. Following fine-tuning, we calculated contextual transcriptional similarity (CTS) for gene pairs from fine-tuned gene embeddings and quantified overall CTS by computing the area under the curve (AUC) for TAD and non-TAD gene pairs (**Fig. 5F**).

To assess developmental changes in transcriptional organization, we calculated the difference in overall AUC (averaged across TAD and non-TAD gene pairs) between Early and Mature stages for each cell type. This analysis allowed us to identify which cell types exhibited the greatest change in AUC as they transitioned from Early to Mature stages, providing insights into the dynamics of transcriptional regulation during cellular maturation (**Fig. 5G**).

### Transcriptional clustering regarding TAD usage in Cancer

For TAD usage in cancer, we compare bulk RNA-seq measurements of normal and primary-tumor tissue in blood, brain, lung, and renal cancers from The Cancer Genome Atlas (TCGA) database^93^. As a caveat, we note that our analysis can only infer correlation, not causation: while mutations leading to the mis-specification of TAD boundaries have been associated with certain cancers^121–123^, there are diverse epigenetic mechanisms underpinning tumorigenesis^124–126^, and the increased expression clustering we observe in the TADs of tumor cells could either be a cause or an effect of these mechanisms. Another caveat regarding our analysis is that the gene groupings were inferred from the consensus TAD scaffold, but in some cancer cells the TAD architecture may have changed. However, the statistical result that these gene groups are over-represented in tumor cells’ transcriptional profiles nonetheless remains valid.

### CTS in cancer datasets

To assess how cancerous states and treatment conditions affect contextual transcriptional similarity (CTS), we analyzed scRNA-seq data from a recent colorectal cancer dataset by Moorman et al.^94^, which tracks cellular plasticity during metastasis (data available at https://github.com/dpeerlab/progressive-plasticity-crc-metastasis). The dataset includes non-cancerous cells, primary cancer cells, and metastatic cancer cells, with each cell type categorized further by treatment status—either treated with 5-fluorouracil-based chemotherapy or untreated. This totals six distinct datasets.

For each dataset, we fine-tuned the pre-trained scGPT model for 10 epochs, using the same settings as described for cell type-specific immune datasets. Following fine-tuning, we calculated CTS for gene pairs and quantified overall CTS by computing the area under the curve (AUC) for both TAD and non-TAD gene pairs. This analysis enabled a systematic comparison of transcriptional dynamics across cancerous and non-cancerous states, as well as between treated and untreated conditions.

### TAD signatures: probabilistic model

We define a 2-component mixture model to infer TAD activation probabilities of a single-cell dataset. Let ***X*** ∈ ℝ^*n*×*p*^ be the gene expression values for a single-cell dataset with *n* cells and *p* genes, with a particular cell *c*’s expression being ***x***^(*c*)^ ∈ R^*p*^. We assume that transcriptionally active TADs (“ON”) correspond to a higher rate of per-gene expression while inactive TADs (”OFF”) correspond to lower rates of gene expression. As mentioned earlier, our model allows inactive TADs to also generate non-zero gene expression, albeit at a lower rate than the “active” TADs. Doing so increases our robustness to noise and allows for one-off gene expression in a TAD. In the mixture model, the probability of ***x***^(*c*)^’s expression in *c* is

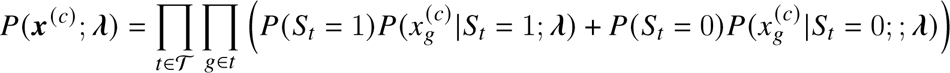

where ***λ*** = {*λ*_*ON*_, *λ*_*OFF*_ }; 𝒯 is the set of all TADs with one or more genes; each TAD *t* ∈ 𝒯 is a set of genes, with *g* being one such gene; *S*_*t*_ is the Bernoulli random variable indicating the activation state of *t*, with *S*_*t*_ = 1 or 0 corresponding to *t* being “ON” or “OFF”, respectively; and 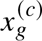 is the expression of gene *g* in the cell *c*. Here, 𝒯, *t* and the gene memberships in TADs are sourced from the species-specific TAD Map. We model that gene expression values are Poisson-distributed counts, though this assumption can be relaxed:

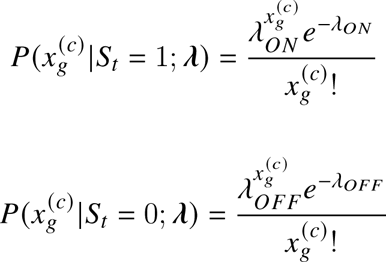

To infer the TAD signature for a particular dataset, we fit this model with the expectation maximization (EM) algorithm, seeking to maximize the log-likelihood over all cells:

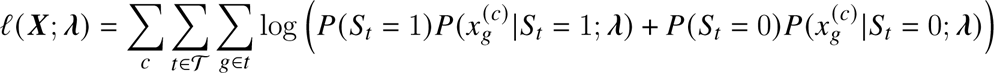

Since the maximum likelihood estimator of the Poisson rate parameter is just the sample mean, the implementation of the EM algorithm is simplified: each maximization round assigns *λ*_*ON*_ and *λ*_*OFF*_ as averages of observed gene expression values across all genes, weighted by the containing TAD’s activation probability *P*(*t* = 1). Also, we note that quasi-Poisson generalizations result in the same maximum likelihood estimator for the expected value of the rate parameter, suggesting that even in cases where the Poisson assumptions do not hold, the corresponding estimate is reasonable.

### Log-odds transformation of TAD signatures

We recommend a log-odds transformation when using TAD signatures to generate data representations for clustering, visualization, or predictive analysis. Many such analyses implicitly or explicitly rely on Euclidean distances between observations. The log-odds transformation converts probabilities (which are in [0, 1] range) to the full range of values in R, making it more amenable to such distance measures.

### TAD signatures for cancer cell line heterogeneity

We acquired scRNA-seq data for cancer cell lines from Kinker et al.’s pan-cancer study^127^, limiting ourselves to cell lines for which drug response data from the PRISM study was also available^128^; this resulted in data on 51,321 cells spanning 193 cancer cell lines. For each cell line, we computed six population statistics: mean, standard deviation and skew, based on either TAD signatures or log-and-count-normalized transcript counts. We reduced each cell line’s drug repsonse profile to the top 50 PCs and considering each PC as an independent regression target, we performed a principal component regression using per-cell-line statistics. Thus a total of 300 (50 × 6) regressions were performed, each with 193 observations. In each regression, the **x** values were reduced to the top 10 principal components, ensuring that all regressions had identical complexity. An *R*^2^ was computed for each regression, indicating the predictive power of the scRNA-seq summary statistic against the drug response measure.

### TAD signatures for cell type inference

We acquired single-cell data from the https://cellxgene.cziscience.com/ portal, obtaining *AnnData*-formatted^129^ datasets from single-cell RNA-seq studies of the human lung (10x sub-study; European Genome-Phenome Archive accession EGAS00001004344; 65,662 cells;^130^), T-cells (GEO accession GSE126030; 51,876 cells;^131^), and breast epithelial cells (GEO accession GSE164898; 31,696 cells;^132^). Cells with fewer than 20 active genes and genes active in less than 10 cells were removed. For the breast tissue data, the dataset annotations seemed to suggest samples were grouped in two broad batches and, to reduce batch effects, we limited ourselves to the larger batch (17,153 cells). The data was then count normalized (to 10^6^) and log transformed using *Scanpy*. Gene identifiers were converted to Ensemble v102, genes were mapped to the human TAD Map inferred in this work, and TAD signatures were estimated. For each dataset, we then generated the following representations: i) principal component analysis (PCA) with 50 components, ii) log-odds (= log 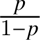) transformation of the TAD signatures computed on the dataset, followed by a 50-component PCA, and iii) a concatenation of the previous two representations. Leiden clustering^133^ using *Scanpy* was performed on each of these representations. The datasets from cellxgene portal contained expert-annotated cell-type labels for each cell and we computed the adjusted Rand index (ARI) of the overlap between computed Leiden clusterings and the expert labels.

#### Quantification and Statistical Analysis

Statistical tests were conducted using version 1.3.1 of the SciPy Python package and R version 4.1.1.

#### Software availability, utility, and efficiency

Pre-computed TAD Maps and consensus TAD boundary estimates for human and mouse genomes are available at http://singhlab.net/tadmap. TAD signatures can be computed using the Python package tadmap, available via pip, conda or GitHub. The package also provides direct programmatic access to the TAD Map. Documentation for the package is available at https://tadmap.readthedocs.io/. TAD signature computations do not require a GPU and can be performed on a personal computer: processing of a scRNA-seq dataset comprising 51,321 cells required 7 minutes of run-time and 8 GB of memory when using a single Intel Xeon 3.47 GHz processor.

## Acknowledgements

We thank Anders Sejr Hansen and Brian Hie for helpful discussions. RS and BB were partially supported by the NIH grants R01GM081871 and R35GM141861. RS and HL were also supported by the Chan-Zuckerberg Initiative.

## Author Contributions

All authors conceived of, contributed to, and wrote the paper. RS developed the TAD Map method with BB; HL led the scGPT integration and the related analysis.

## Supplementary Materials

**Figure S1:**
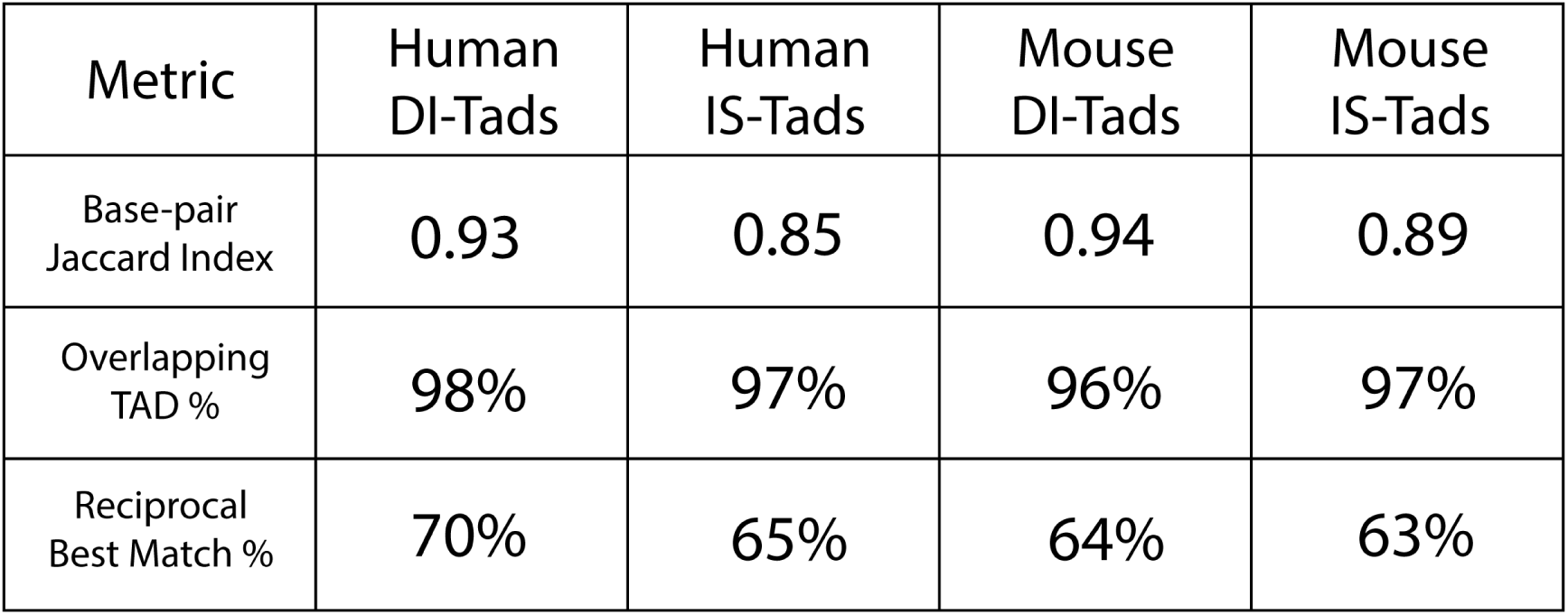
TAD Maps with expanded cell types and TAD-calling algorithms strongly agree with originals. Base-pair Jaccard Index measures nucleotide-level agreement between expanded and original TAD maps as the ratio of overlapping TAD base pairs to their union; TAD-level overlap reflects the fraction of expanded TADs overlapping any original TAD; reciprocal best matches indicates one-to-one correspondence using a greedy one-to-one matching scheme based on genomic distance, which enforces unique pairings between TADs.

**Figure S2:**
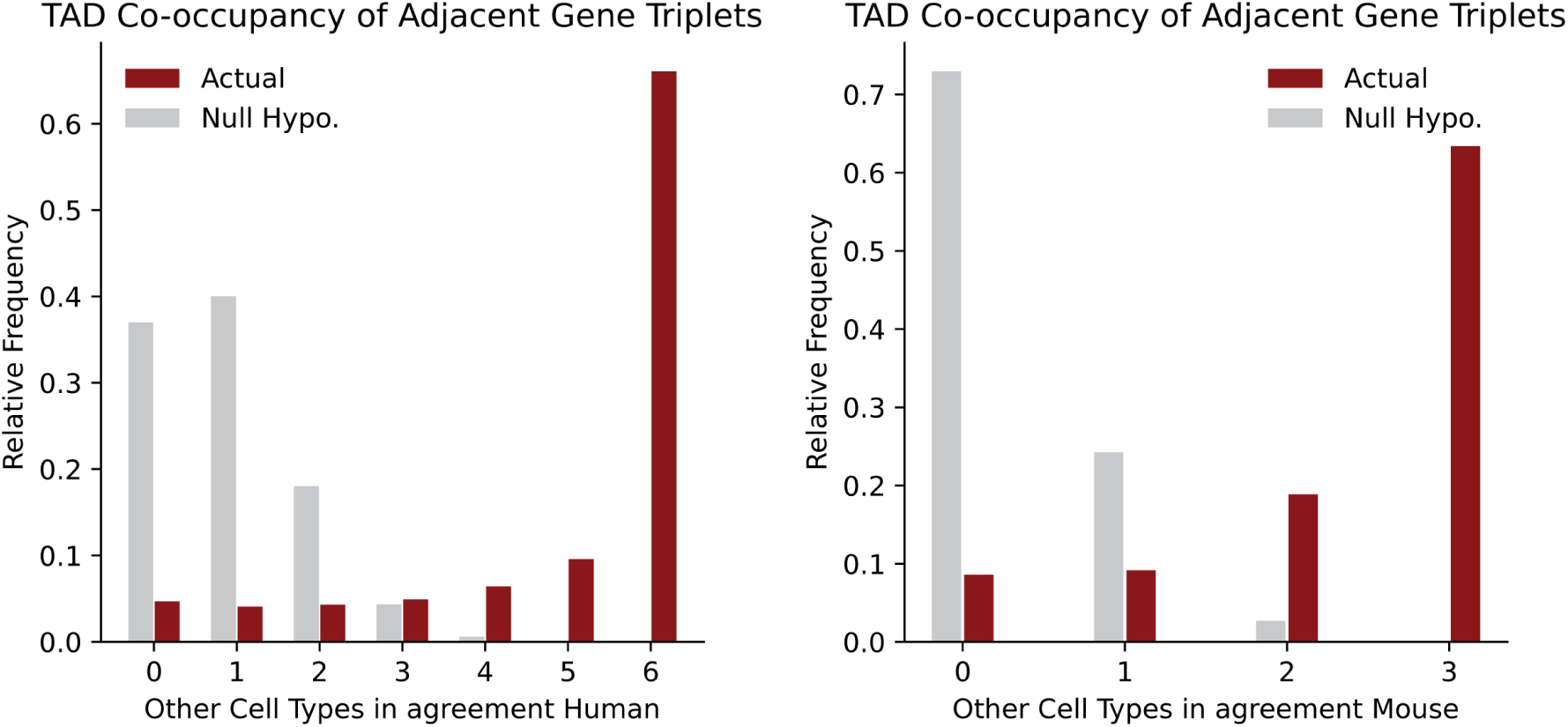
Adjacent gene triplets yield consistent results with gene pairs. Like pairs, triplets that co-occupy the same TAD across all cell types exceed expectations under a null model. This figure complements Fig. 2C.

**Figure S3:**
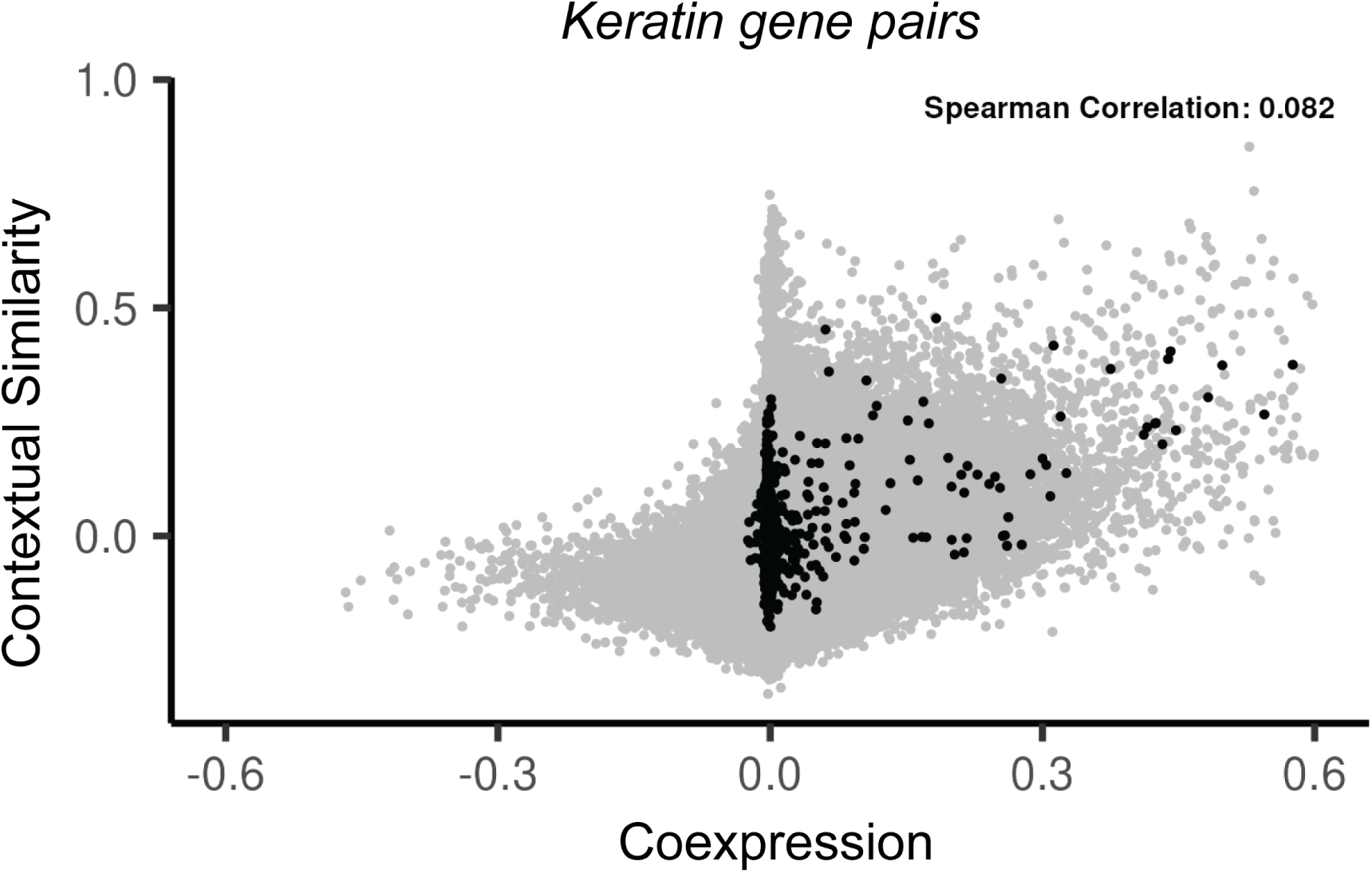
Contextual similarities (CTS) vs co-expression for keratin genes pairs. This figure complements Fig. 3E-G, where it visualizes CTS and co-expression for all gene pairs in the keratin family.

**Figure S4:**
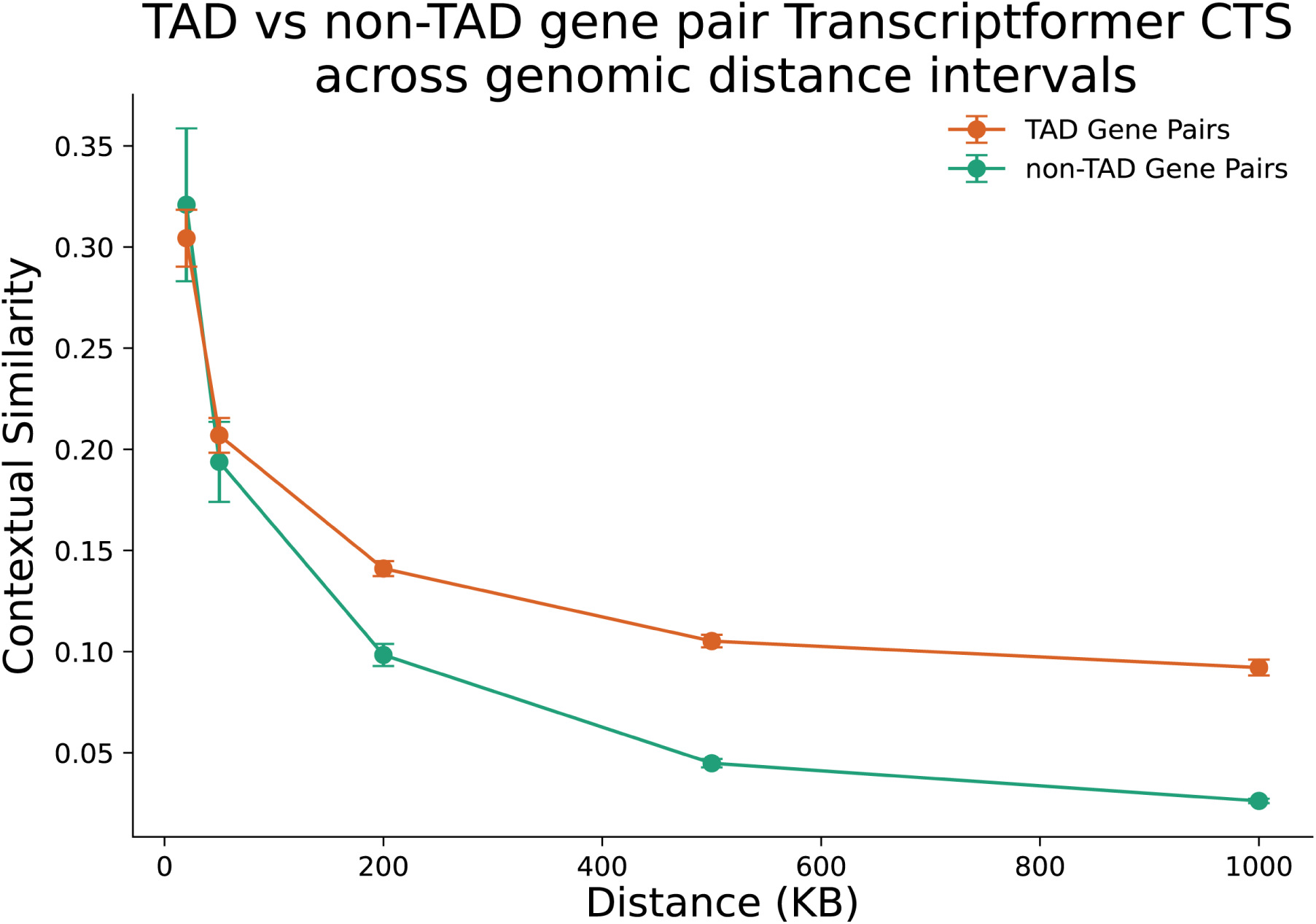
Contextual similarities (CTS) derived from Transcriptformer show consistent, if not stronger, patterns compared to scGPT CTS across the same genomic distance intervals. This figure complements Fig. 4B.

**Figure S5:**
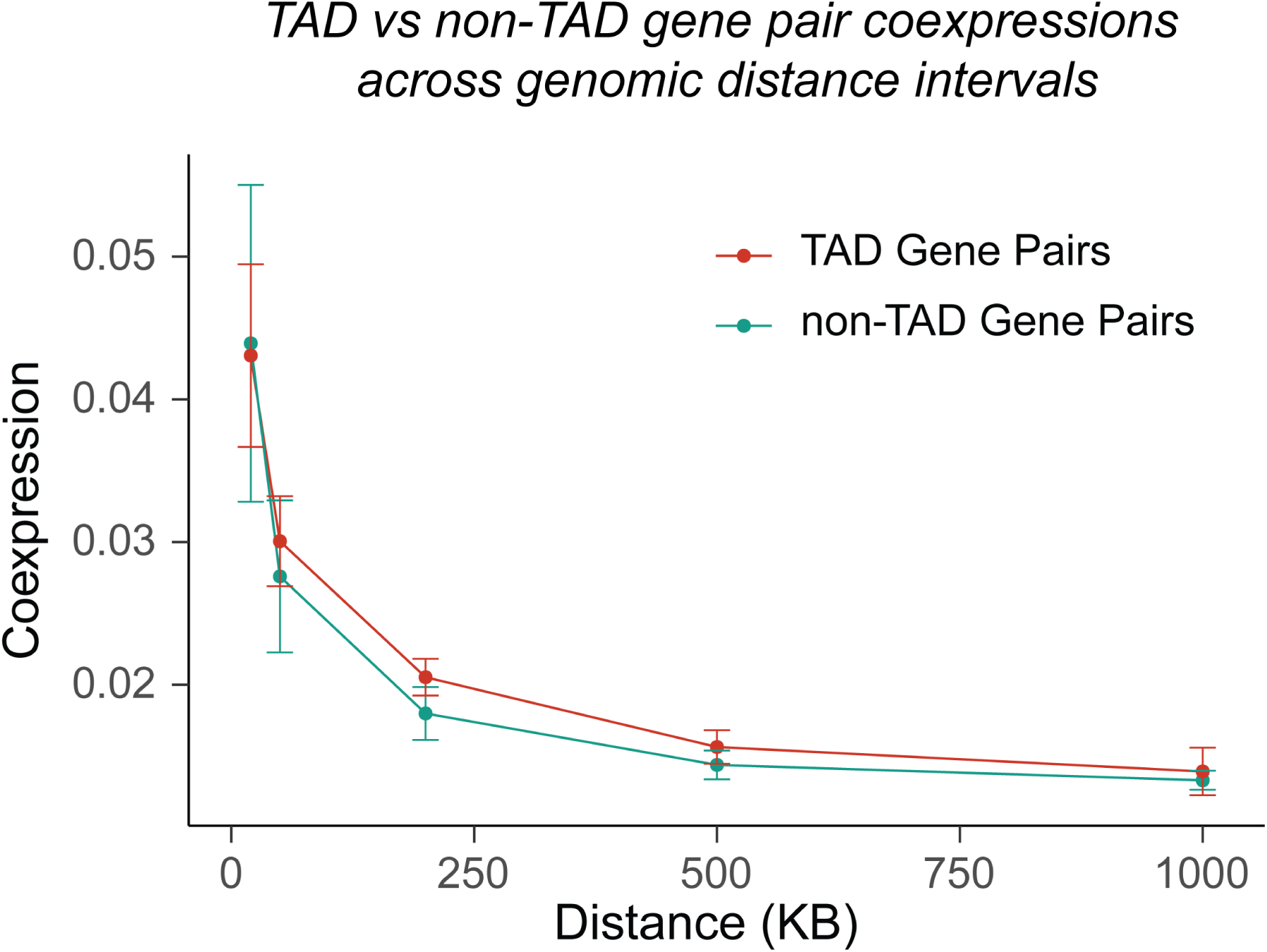
Gene pair co-expression exhibits inconsistent patterns with respect to distance. This figure contrasts the analysis with CTS in Fig. 4B, highlighting that average co-expression shows irregular and inconsistent trends between TAD and non-TAD gene pairs across distance intervals.

**Figure S6:**
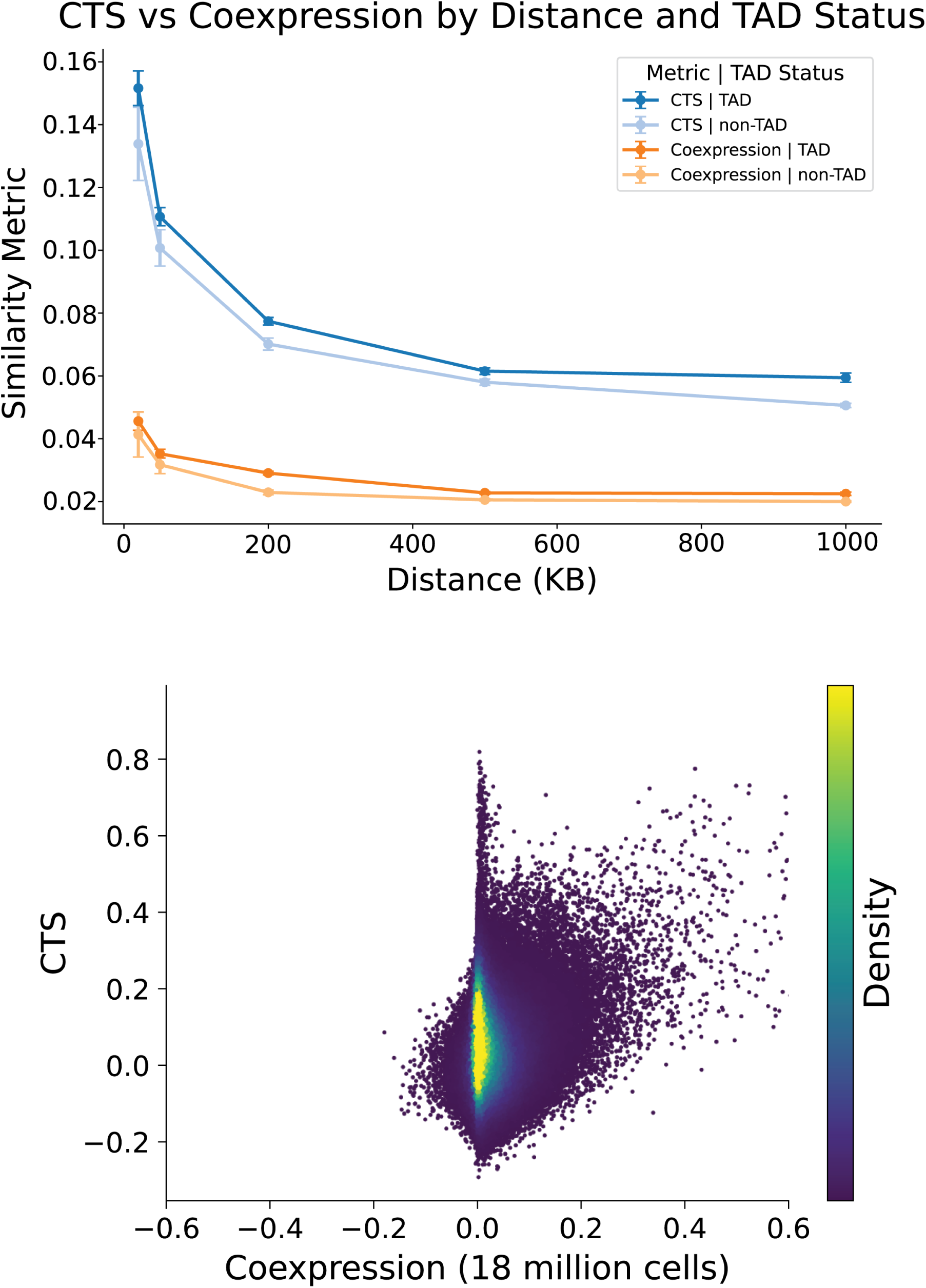
Genomic distance–binned and unbinned co-expression analyses across 700 additional single-cell datasets from scBaseCount yield patterns consistent with the primary co-expression results. The top figure complements Fig. S5 and the bottom figure complements Fig. 3D, extending both analyses to 18 million cells across 40 tissues.

**Figure S7:**
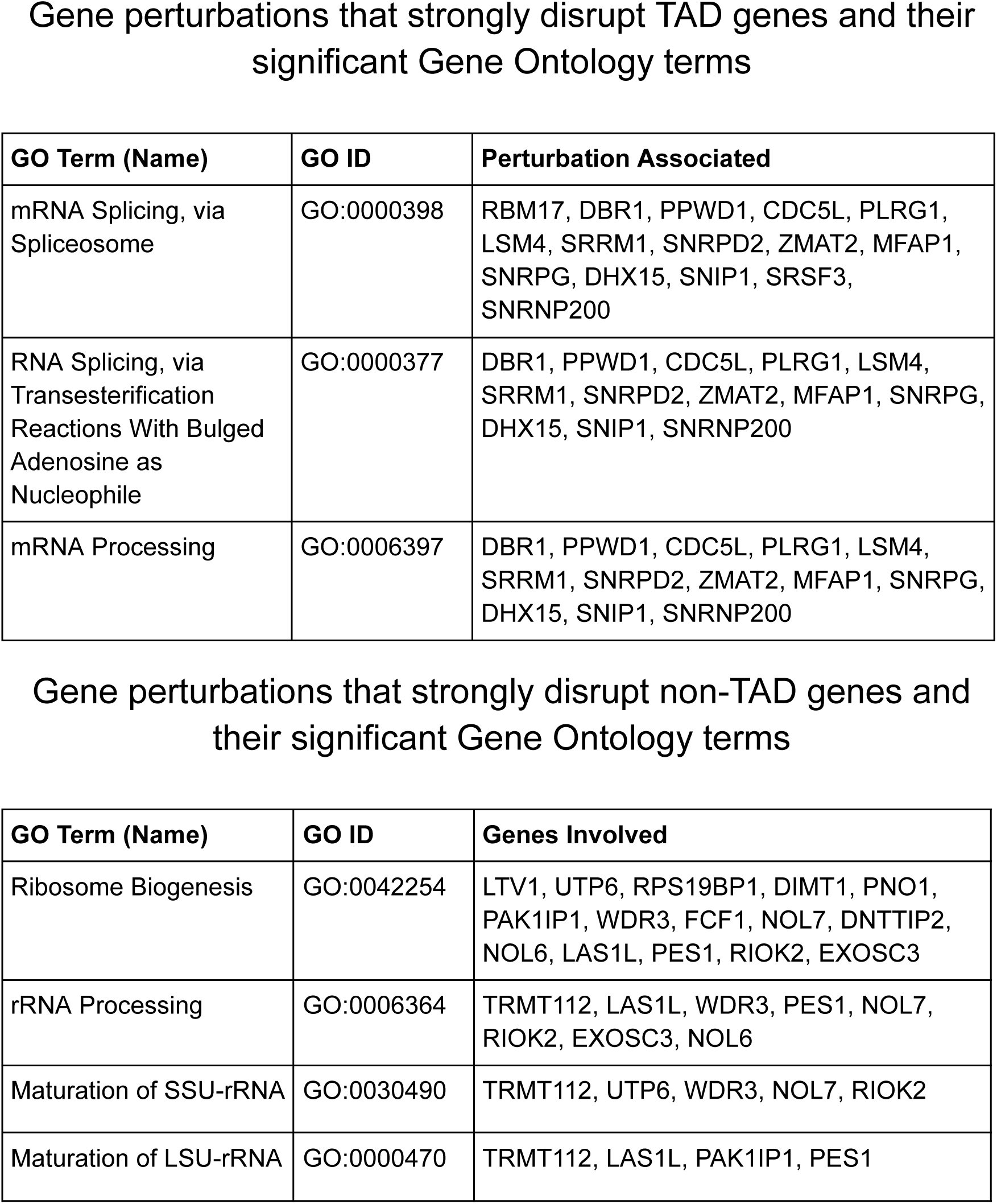
Extended Perturb-seq analysis with additional datasets to assess perturbed genes that most disrupt transcription in TAD and non-TAD regions. The top panel mirrors the original analysis, highlighting genes involved in transcriptional regulation and mRNA processing (complementing Fig. 4G), while the bottom panel similarly highlights genes associated with ribosomal components and rRNA processing (complementing Fig. S8D).

**Figure S8:**
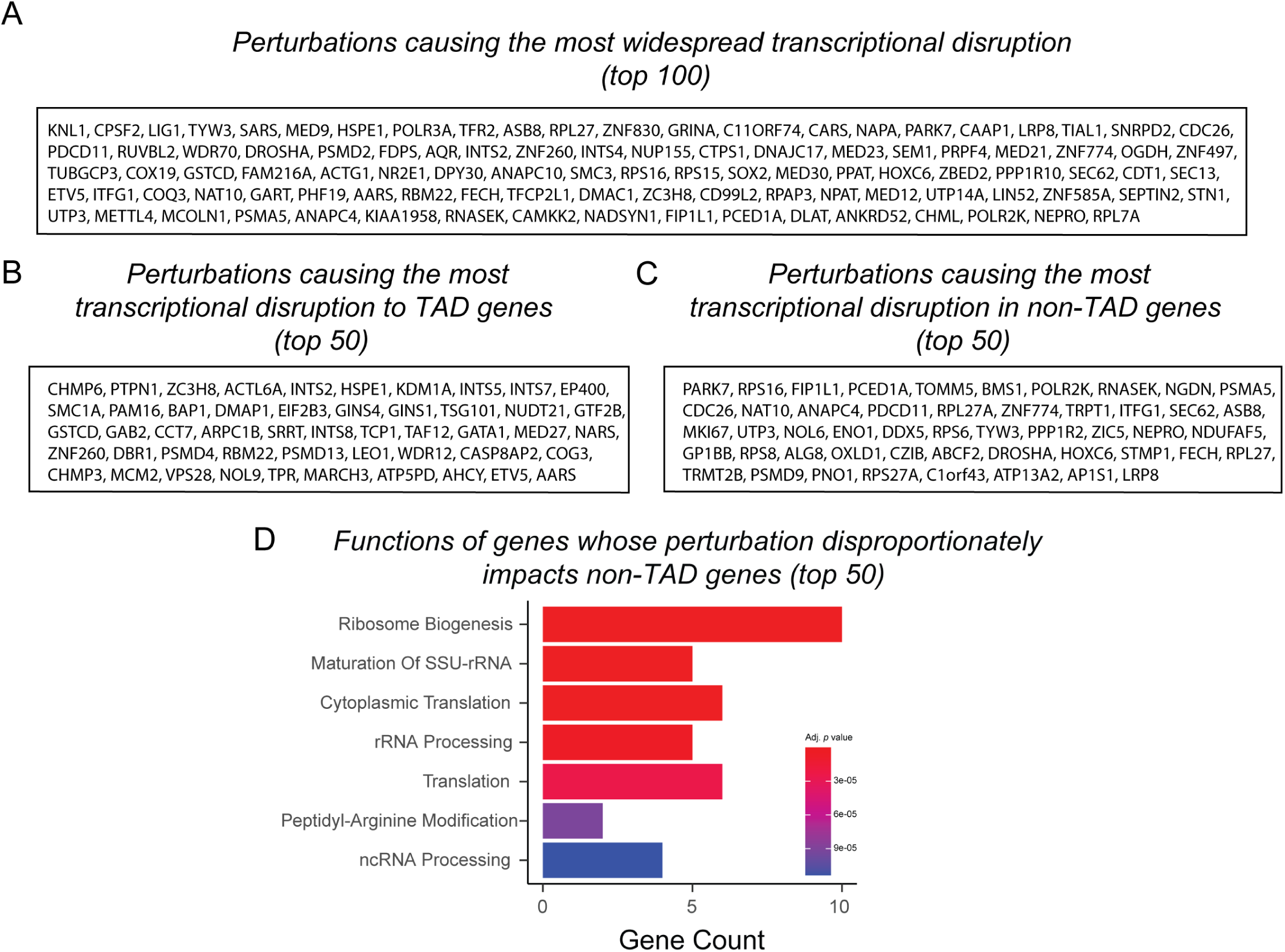
List of specific genes whose perturbation induces transcriptional disruption across TAD and non-TAD regions. **A**) This gene group highlights the genes whose perturbation causes the most widespread transcriptional disruption overall, without distinguishing between TAD and non-TAD regions. **B**) This gene group highlights genes whose knockdown disproportionally disrupts transcription within TADs. Corresponding Enrichr analysis of this gene set is shown in Fig. 4G. **C**) This gene group highlights genes whose knockdown leads to the greatest influence on transcriptional disruption specifically within TADs. **D**) This panel displays the Enrichr analysis of genes whose knockdown disproportionately disrupts transcription in non-TAD regions.

**Figure S9:**
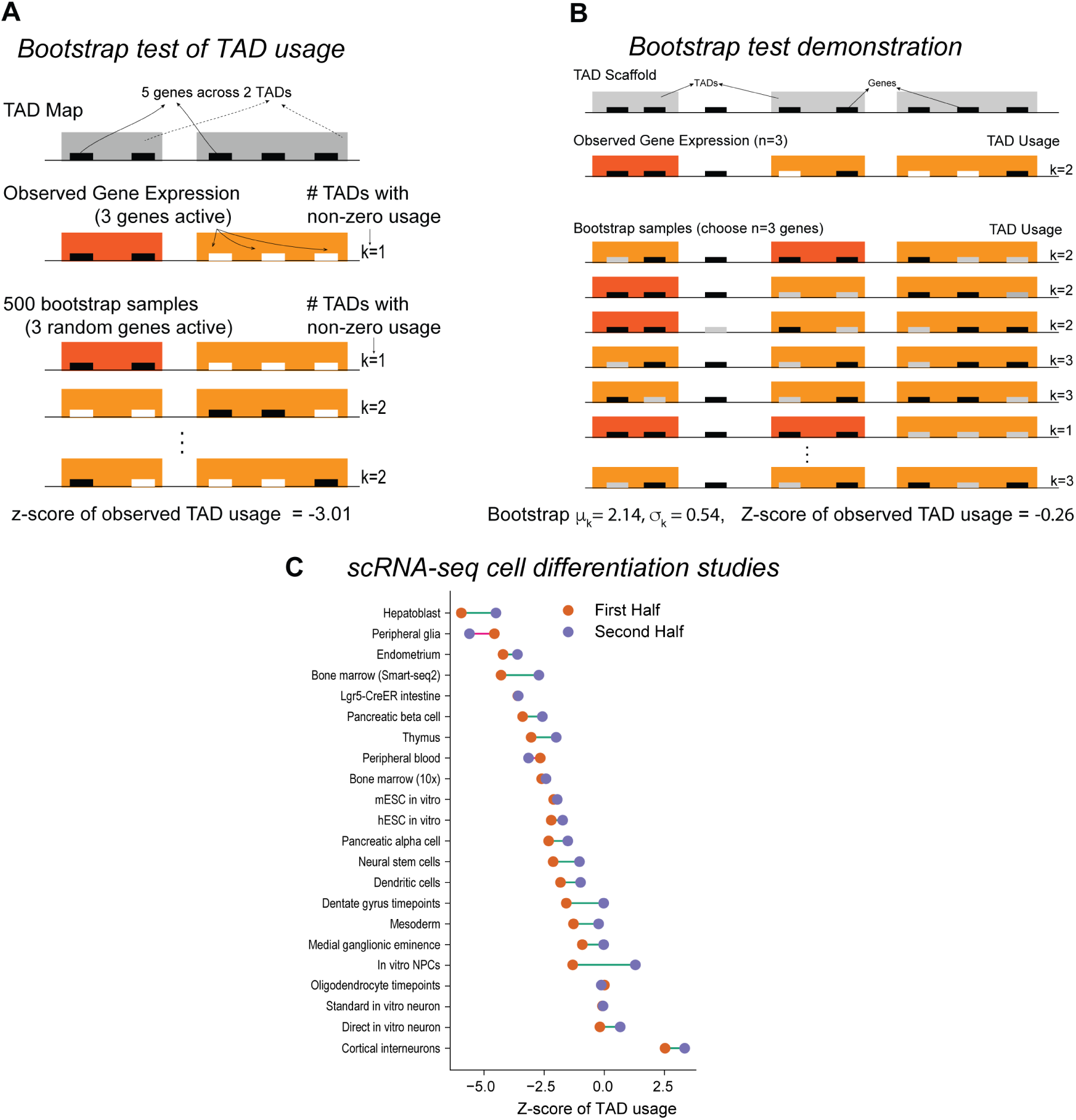
Bootstrap test to assess gene clustering in TADs. **A**) We define a TAD’s *usage* as the number of transcriptionally active genes within it. Our bootstrap test assumes gene activity is independent of TADs under the null hypothesis. For any transcriptional profile, we randomize gene activity while preserving the total number of active genes, expressing TAD usage as a z-score relative to the expected distribution. This approach is robust to noise, sparsity, and platform effects, particularly in scRNA-seq data. Bootstrap z-scores can be aggregated across datasets or used to distinguish highly versus moderately active TADs. **B**) The bootstrap test is demonstrated here with a more complex artificial example. **C**) Gene clustering into TADs decreases during cell differentiation, as shown in Figure 5A. Early differentiation stages display stronger clustering of genes into TADs. In Figure 5A, cells were divided into “first stage vs. the rest” based on the CytoTrace Order variable**^?^**. Here, cells are partitioned into two groups with equal sizes. Regardless of partitioning, the clustering trend remains consistent.

**Figure S10:**
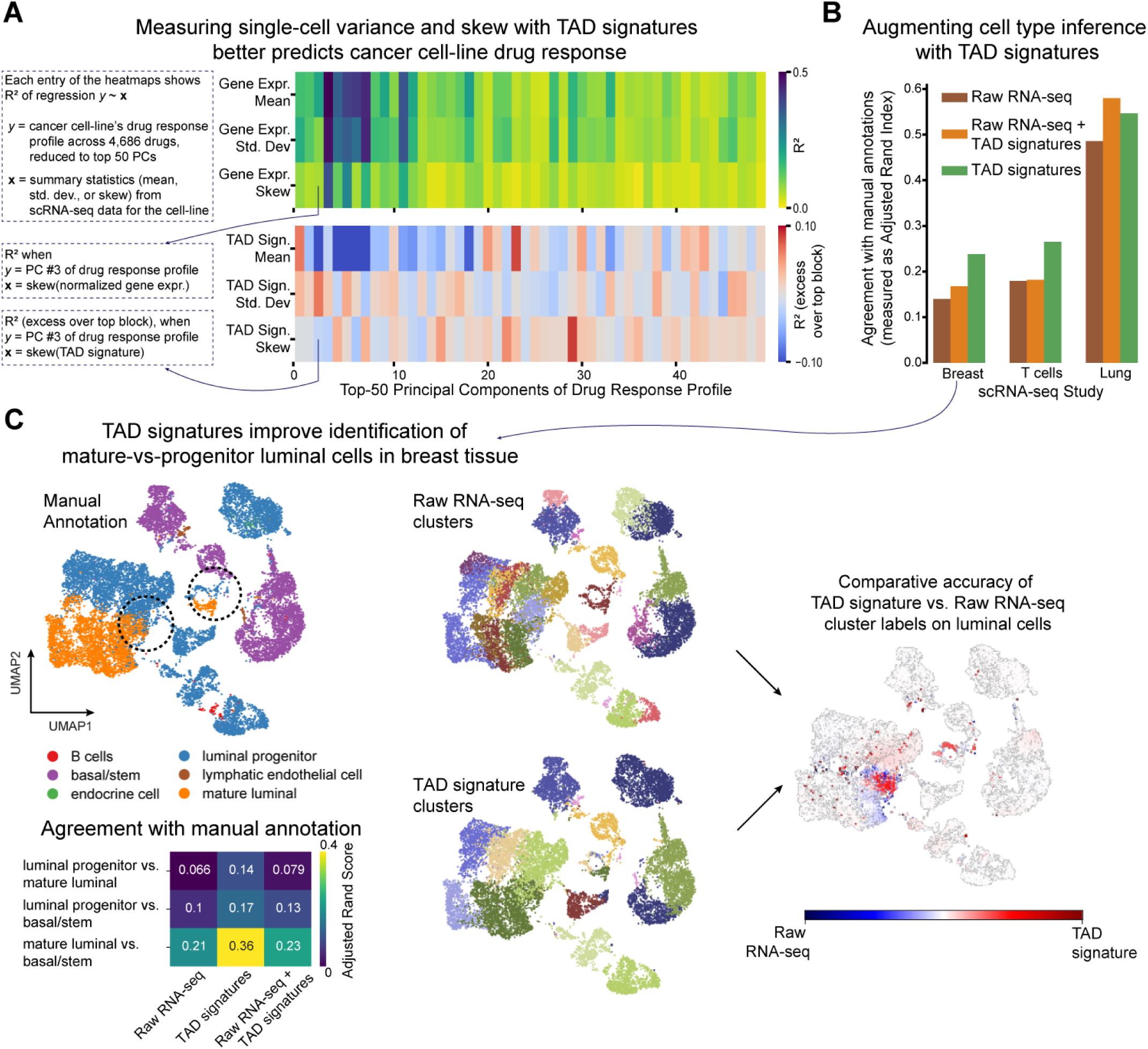
TAD signatures augment standard scRNA-seq analysis. **A**) TAD signatures reveal biologically meaningful single-cell heterogeneity. For 193 cancer cell lines, we analyzed drug response profiles from the PRISM study^128^, reduced to 50 principal components. We computed the first three moments (mean, standard deviation, skew) of scRNA-seq data, either as log-normalized gene expression (top block) or log-odds of TAD signatures (bottom block). The second and third moments capture within-line heterogeneity. We used principal component regression (*k* = 10) to predict drug responses from per-gene or per-TAD statistics and measured predictive power as *R*^2^. For the second and third moments, TAD signatures were significantly more predictive than gene expression (e.g., skew: *p* = 0.0014, one-sided Wilcoxon rank-sum test).**B**) TAD signatures improve automated cell-type inference. Compared to Leiden clusters from scRNA-seq alone, clusters using only TAD signatures or combining TAD and scRNA-seq better matched manual cell-type annotations in three scRNA-seq datasets. For breast and T cell data, TAD signatures alone outperformed combined clustering, likely due to RNA-seq noise. **C**) In breast tissue data, TAD signatures more accurately distinguished progenitor from mature luminal cells, highlighting their value in studying cell differentiation. Middle-column plots show Leiden clusters; RNA-seq clustering produced more clusters, aligning less with biological distinctions.

